# Oncogenic Kras^G12D^ specific non-covalent inhibitor reprograms tumor microenvironment to prevent and reverse early pre-neoplastic pancreatic lesions and in combination with immunotherapy regresses advanced PDAC in a CD8^+^ T cells dependent manner

**DOI:** 10.1101/2023.02.15.528757

**Authors:** Krishnan K. Mahadevan, Kathleen M. McAndrews, Valerie S. LeBleu, Sujuan Yang, Hengyu Lyu, Bingrui Li, Amari M. Sockwell, Michelle L. Kirtley, Sami J. Morse, Barbara A. Moreno Diaz, Michael P. Kim, Ningping Feng, Anastasia M. Lopez, Paola A. Guerrero, Hikaru Sugimoto, Kent A. Arian, Haoqiang Ying, Yasaman Barekatain, Patience J. Kelly, Anirban Maitra, Timothy P. Heffernan, Raghu Kalluri

## Abstract

Pancreatic ductal adenocarcinoma (PDAC) is associated with mutations in Kras, a known oncogenic driver of PDAC; and the *KRAS^G12D^* mutation is present in nearly half of PDAC patients. Recently, a non-covalent small molecule inhibitor (MRTX1133) was identified with specificity to the Kras^G12D^ mutant protein. Here we explore the impact of Kras^G12D^ inhibition by MRTX1133 on advanced PDAC and its influence on the tumor microenvironment. Employing different orthotopic xenograft and syngeneic tumor models, eight different PDXs, and two different autochthonous genetic models, we demonstrate that MRTX1133 reverses early PDAC growth, increases intratumoral CD8^+^ effector T cells, decreases myeloid infiltration, and reprograms cancer associated fibroblasts. Autochthonous genetic mouse models treated with MRTX1133 leads to regression of both established PanINs and advanced PDAC. Regression of advanced PDAC requires CD8^+^ T cells and immune checkpoint blockade therapy (iCBT) synergizes with MRTX1133 to eradicate PDAC and prolong overall survival. Mechanistically, inhibition of mutant Kras in advanced PDAC and human patient derived organoids (PDOs) induces Fas expression in cancer cells and facilitates CD8^+^T cell mediated death. These results demonstrate the efficacy of MRTX1133 in different mouse models of PDAC associated with reprogramming of stromal fibroblasts and a dependency on CD8^+^ T cell mediated tumor clearance. Collectively, this study provides a rationale for a synergistic combination of MRTX1133 with iCBT in clinical trials.

## Introduction

*KRAS* mutations are a dominant genetic event in pancreatic adenocarcinoma (PDAC), with approximately 90% of patients presenting with such mutations, and 45% of patients having *KRAS^G12D^* mutations (Raphael et al., 2017). Genetically engineered mouse models revealed that Kras^G12D^ is critical for the initiation and maintenance of PDAC (Collins et al., 2012; Hingorani et al., 2003; Hingorani et al., 2005; Ying et al., 2012), suggesting that mutant Kras is a viable therapeutic target for the control of PDAC progression. Recently, inhibitors targeting the switch II pocket of mutant Kras^G12C^ and Kras^G12D^ have been developed, enabling suppression of mutant Kras and downstream RAF-MEK-ERK (MAPK) signaling (Hallin et al., 2022; Hallin et al., 2020; Lanman et al., 2020; Ostrem et al., 2013; Wang et al., 2022); however, the precise underlying mechanisms for the efficacy of such inhibitors are not fully understood. Mutant Kras promotes tumor initiation and progression through a number of cell intrinsic mechanisms, including promoting cancer cell proliferation and rewiring cancer cell metabolic dependencies to promote growth (Commisso et al., 2013; Dey et al., 2020; Yao et al., 2019; Ying et al., 2012). In addition, mutant Kras can act through cancer cell extrinsic mechanisms to remodel the tumor microenvironment and impact cancer initiation and progression (Pylayeva-Gupta et al., 2011).

The complex desmoplastic response associated with PDAC is characterized by infiltration of immune cells and accumulation of fibroblasts and extracellular matrix. Studies with defined mutational drivers in cancer cells support a role of mutant Kras in remodeling the tumor immune microenvironment (TIM) and, reciprocally, indicate that the TIM regulates the initiation and progression of cancer. Specifically, CD11b^+^ myeloid cells promote the initiation and maintenance of Kras^G12D^ driven pancreatic cancer (Zhang et al., 2017). Kras^G12D^ is also associated with increased infiltration of CD4^+^ T cells (Brembeck et al., 2003) and mutant Kras^G12V^ promotes the induction of Tregs (Zdanov et al., 2016), potentially contributing to the increase in Treg abundance during PDAC progression (Hiraoka et al., 2006). Lung cancer and preinvasive pancreatic neoplasia driven by Kras^G12D^ are associated with infiltration of IL17^+^CD4^+^ T cells (Th17) cells, which functionally promote tumor initiation and progression (Chang et al., 2014; McAllister et al., 2014). Studies with genetic suppression of *Kras^G12D^* in PDAC revealed that mutant Kras^G12D^ drives the accumulation of immunosuppressive CD4^+^Gata3^+^ Th2 cells through IL-33 secretion by cancer cells (Alam et al., 2022; Dey et al., 2020). Moreover, pharmacological Kras^G12C^ inhibition with AMG510 increased T cell infiltration and was synergistic with immune checkpoint blockade therapy (iCBT) (Canon et al., 2019), further supporting that mutant Kras associated remodeling of the TIM may be leveraged therapeutically to limit cancer progression. PDAC cancer associated fibroblasts (CAFs) are also implicated in cancer progression, with emerging findings on their functional heterogeneity (Chen et al., 2021; McAndrews et al., 2022; Ozdemir et al., 2014; Rhim et al., 2014), though their specific phenotypic distribution in the context of Kras^G12D^ inhibition is unknown.

Here, we evaluate the efficacy of Kras^G12D^ (Kras*) inhibition with the small molecule inhibitor MRTX1133 in orthotopic and spontaneous mouse models of PDAC. Kras* inhibition was associated with TIM remodeling and reprogramming of CAFs. We demonstrate that cancer cell clearance upon treatment of advanced PDAC with MRTX1133 or by *in-vivo* RNA interference targeting Kras^G12D^ (siKras^G12D^ electroporated in exosomes) is mediated by Fas-FasL interactions between cancer cells and CD8^+^ T cells, and that iCBT synergizes with MRTX1133 to eradicate advanced PDAC, providing a rationale for clinical testing of such combination therapy.

## Results

### Kras* inhibition suppresses the growth of human PDAC cells

To evaluate the impact of Kras* inhibition on pancreatic cancer cells, a panel of cell lines were treated with MRTX1133. Treatment of a human PDAC *KRAS^G12D^* mutant cell line (HPAC) and a murine PDAC *Kras^G12D^* mutant cell line (KPC689) with MRTX1133 resulted in effective downregulation of pERK (**Fig. 1A**), a downstream readout of Kras* activity. In contrast, pERK was not impacted in KPC689 cells treated with the Kras^G12C^ inhibitor MRTX849 (**Supplementary Fig. 1A**). MRTX1133 inhibited the proliferation of cell lines with Kras* mutation but had reduced impact on cell lines with other Kras mutations (**Fig. 1B**, **Supplemental Fig. 1B**). Collectively these results support the specific efficacy of MRTX1133 for PDAC cells with Kras*.

**Figure 1:**
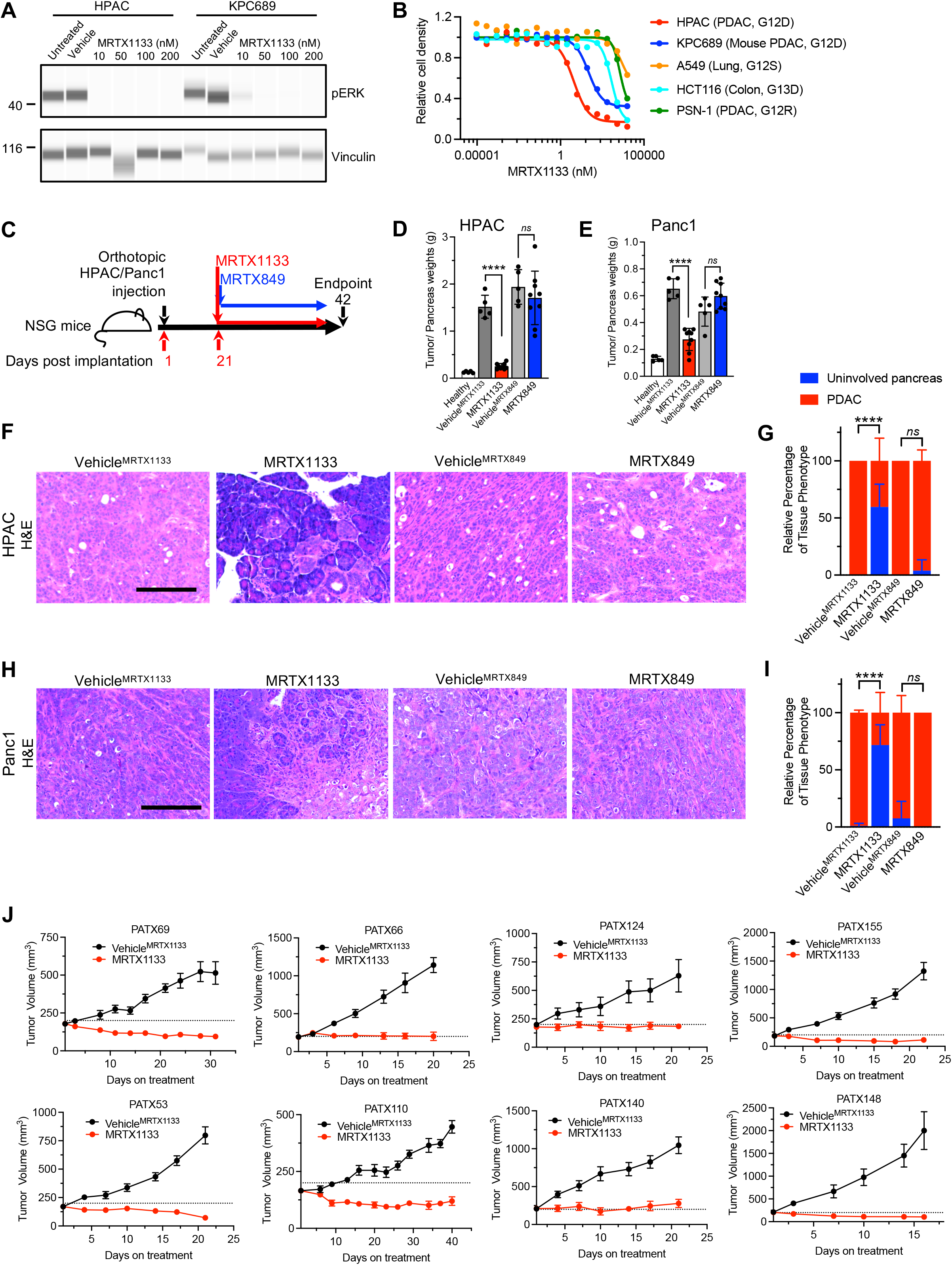
MRTX1133 controls the growth of human cell lines and PDXs. **(A)** Capillary immunoassay for pERK and vinculin abundance in HPAC and KPC689 cells in response to MRTX1133. Vehicle: DMSO. (**B**) Cell proliferation in response to MRTX1133. Cell lines and tissue of origin, and mutational status of *KRAS/Kras* are listed in parentheses (n=3 biological replicates/ group). (**C**) Schematic of orthotopic experiments in NSG mice with HPAC and Panc1. Treatment started at day 21, and mice were euthanized at day 42. (**D-E**) Tumor weights at endpoint for orthotopic HPAC (Healthy: n=5, Vehicle^MRTX1133^: n=5, MRTX1133: n=9, Vehicle^MRTX849^: n=5, MRTX849: n=9) (**D**) and Panc1 (Healthy: n=5, Vehicle^MRTX1133^: n=5, MRTX1133: n=9, Vehicle^MRTX849^: n=5, MRTX849: n=9) (**E**). (**F-I**) Histological analyses of orthotopic HPAC (Vehicle^MRTX1133^: n=4, MRTX1133: n=5, Vehicle^MRTX849^: n=4, MRTX849: n=6) and Panc1 (Vehicle^MRTX1133^: n=5, MRTX1133: n=4, Vehicle^MRTX849^: n=4, MRTX849: n=5) tumors. (**F**) Representative H&E images of orthotopic HPAC tumors with the indicated treatments. (**G**) Histological scoring of orthotopic HPAC tumors. (**H**) Representative H&E images of orthotopic Panc1 tumors with the indicated treatments. (**I**) Histological scoring of orthotopic Panc1 tumors. (**J**) Tumor growth curves of PDXs implanted subcutaneously into athymic nude mice (n=5-6 replicates per group). In **D, E, G, I** and **J** data are presented as mean + SD. Significance was determined by unpaired-T test in **D, E** and by two-way ANOVA with Sidak’s multiple comparisons test in **G** and **I**. **** P<0.0001, *ns*: not significant. Scale bars: 100 μm.

The anti-tumor efficacy of MRTX1133 on tumor growth kinetics was confirmed using orthotopic implantation of human PDAC cells in NSG mice (**Fig. 1C**). HPAC and Panc1 orthotopic xenografts treated with MRTX1133 (30 mg/kg BID i.p.) showed significantly reduced tumor/pancreas weights, whereas MRTX849 treatment (30 mg/kg QD p.o.) had no impact on tumor growth (**Fig. 1D-E, Supplemental Fig. 1C-D**). The observed reduction in tumor weight was consistent with histological findings, with mostly normal parenchyma and significantly reduced PDAC with MRTX1133 treatment compared to controls (**Fig. 1F-I**). MRTX1133 treatment was not associated with specific changes in bodyweight in either tumor models (**Supplemental Fig. 1E-F**). Next, a panel of eight different Kras* PDXs implanted subcutaneously in athymic nude mice were also evaluated for response to MRTX1133. All eight PDXs demonstrated tumor control or slight regression in response to MRTX1133 (**Fig. 1J**). Despite inhibition of tumor growth, hisopathological analysis at end point showed persistence of cancer cells in the MRTX1133 treated PDX bearing mice (**Supplemental Fig. 1G**). Analysis of target engagement and downstream pathways demonstrated effective downregulation of pERK, DUSP6, and SPRY4 (**Fig. 1J, Supplemental Fig. 2A-F**).

### Established PDAC regresses with Kras* targeting by MRTX1133 in immunocompetent mice

To evaluate the efficacy of MRTX1133 in an immunocompetent background, KPC689 cells were orthotopically implanted in C57BL/6J mice (**Fig. 2A**). MRTX1133 treatment significantly regressed PDAC and improved tumor histology without specific impact on body weight (**Fig. 2B-E, Supplemental Fig. 2G, I**). MRTX1133 treated tumors also displayed reduced pERK abundance (**Fig. 2D, F**).

**Figure 2:**
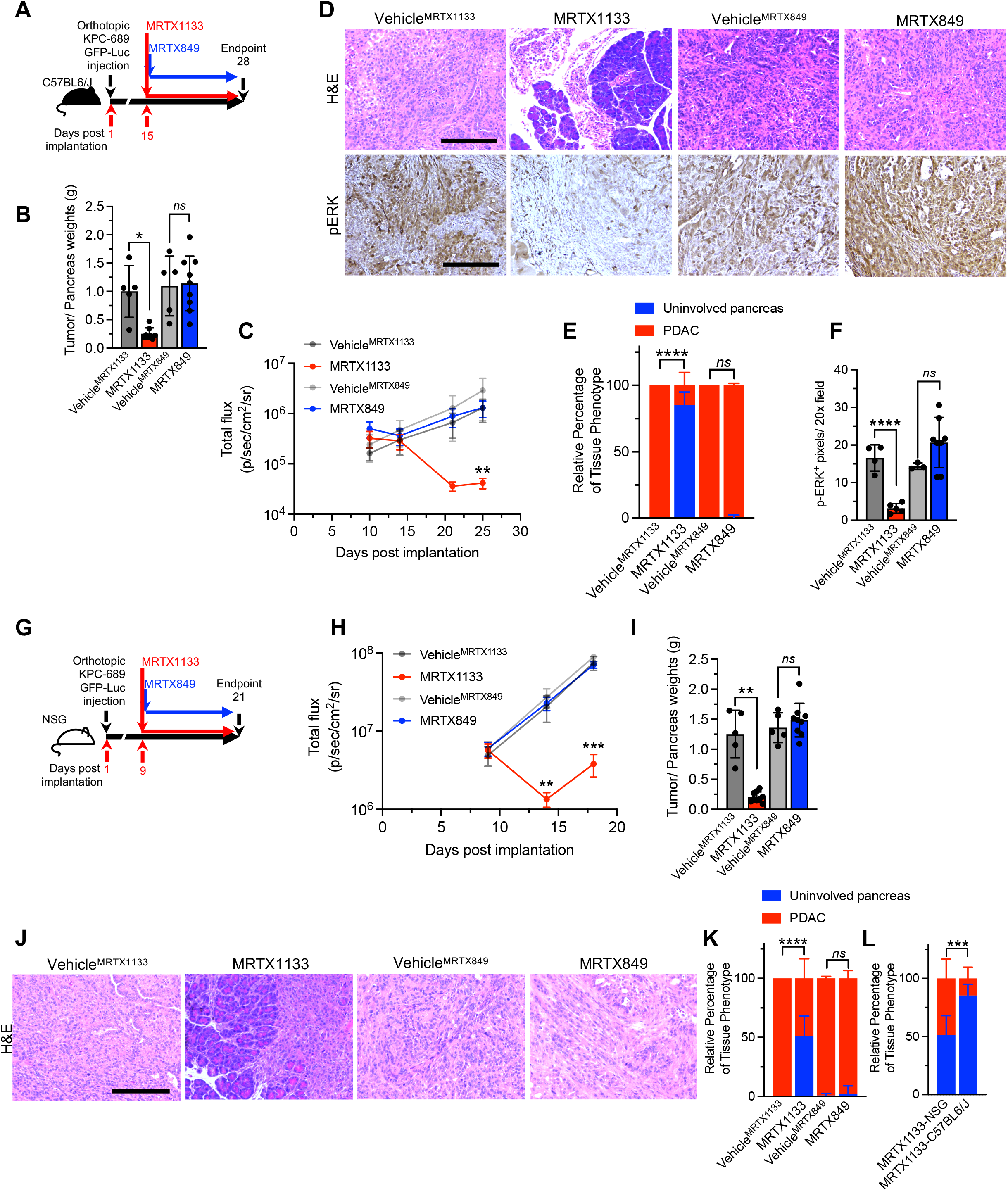
MRTX1133 efficacy in immunocompetent and immunodeficient backgrounds. (**A**) Schematic of experiments with C57BL6/J mice orthotopically implanted with KPC689. Treatment started at day 15, and mice were euthanized at day 28. (**B**) Tumor weights of orthotopic KPC689 tumors in C57BL6/J mice at endpoint (Vehicle^MRTX1133^: n=5, MRTX1133: n=9, Vehicle^MRTX849^: n=5, MRTX849: n=9). (**C**) Bioluminescence of orthotopic KPC689 tumors in C57BL6/J mice over time. Treatment was initiated at day 15 (Vehicle^MRTX1133^: n=5, MRTX1133: n=9, Vehicle^MRTX849^: n=5, MRTX849: n=9). (**D-F**) Representative H&E (Vehicle^MRTX1133^: n=5, MRTX1133: n=6, Vehicle^MRTX849^: n=5, MRTX849: n=8) and pERK (Vehicle^MRTX1133^: n=4, MRTX1133: n=5, Vehicle^MRTX849^: n=3, MRTX849: n=8) immunostaining images of the pancreata of C57BL6/J mice orthotopically implanted C57BL6/J (**D**) and quantification (**E-F).** (**G**) Schematic of experiments with NSG mice orthotopically implanted with KPC689. Treatment started at day 9, and mice were euthanized at day 21. (**H**) Bioluminescence of orthotopic KPC689 tumors in NSG mice over time. Treatment was initiated at day 9 (Vehicle^MRTX1133^: n=5, MRTX1133: n=9, Vehicle^MRTX849^: n=5, MRTX849: n=10) (**I**) Tumor weights of orthotopic KPC689 tumors in NSG mice at endpoint (Vehicle^MRTX1133^: n=5, MRTX1133: n=9, Vehicle^MRTX849^: n=5, MRTX849: n=9) (**J-K**) Representative H&E images of the pancreata of NSG mice orthotopically implanted C57BL6/J (**J**) and quantification (Vehicle^MRTX1133^: n=5, MRTX1133: n=6, Vehicle^MRTX849^: n=4, MRTX849: n=10) (**K**). (**L**) Histological scoring of the pancreata of MRTX1133 treated NSG and C57BL6/J mice orthotopically implanted with KPC689 at endpoint. Data included in this plot are also presented in panel (K) and (E). In **B, C, E, F, H, I, K and L** data are presented as mean + SD. In **B and I**, unpaired -T test was used for comparison of all panels and Welch’s correction was used for vehicle and MRTX1133 comparisons. In **E, K and L**, two-way ANOVA with Tukey’s multiple comparisons test was used. In **C** and **H**, significance was determined by Mann-Whitney test (Vehicle^MRTX1133^ vs. MRTX1133 total flux comparisons). Significance was determined by unpaired-T test in **F**. Scale bar: 100 μm.

In the context of an immunodeficient background lacking T cells, B cells, and NK cells in the NSG mice (**Fig. 2G**), MRTX1133 controlled orthotopic KPC689 tumor growth without specific impact on body weight (**Fig. 2H-K, Supplemental Fig. 2H, J**). Histopathological assessment of MRTX1133 treated tumors showed greater PDAC involvement in NSG mice compared to C57BL/6J mice at end point suggesting that adaptive immune cells may contribute the therapeutic efficacy of MRTX1133 (**Fig. 2L**).

To evaluate the impact of Kras* targeting in the context of tumor initiation and PanIN emergence, genetically engineered mouse models (GEMMs) driven by *Kras^G12D^* expression in the pancreas (KC mice; see methods) were employed. KC mice were treated with MRTX1133 starting at 7 weeks of age, when on average 5% of the pancreas histology involves PanIN lesions (**Fig. 3A-C**). Kras* inhibition marginally decreased pancreas weight (**Supplemental Fig. 3A**) and reduced the percentage of tissue with PanIN from 5% to 0.05% upon 13 weeks of treatment, whereas 35% pancreata of vehicle treated mice exhibited PanIN lesions after 13 weeks (**Fig. 3B-D**). CK19 staining confirmed regressed PanIN lesions in MRTX1133 treated mice (**Fig. 3B, D**). In addition, no specific differences in bodyweight or urine protein levels were observed between vehicle and MRTX1133 treated mice over the course of 13 weeks of treatment (**Supplemental Fig. 3B-C**). Blood biochemistry and histological analyses revealed minimal changes upon 13 weeks of treatment with MRTX1133 compared to controls (**Supplemental Fig. 3D-E**).

**Figure 3:**
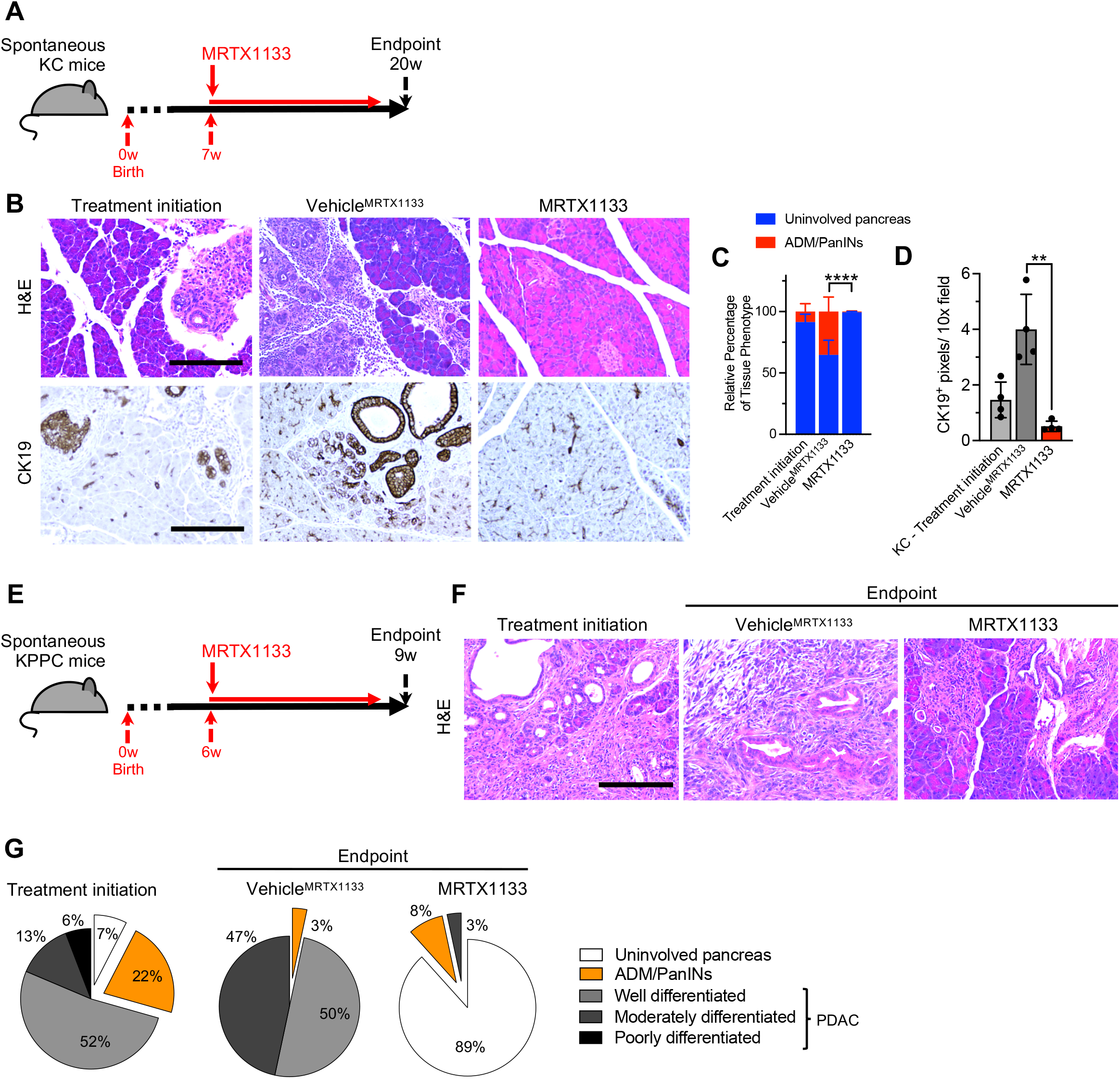
MRTX1133 reverses precancerous and established cancerous lesions. (**A**) Schematic of spontaneous KC treatment experiments. Treatment was initiated at 7 weeks and mice euthanized at 20 weeks post-birth. (**B-D**) Representative H&E and CK19 immunostaining images of the pancreata of KC mice (**B**) and histological (Treatment initiation: n=6, Vehicle^MRTX1133^: n=4 and MRTX1133: n=4) and CK19 immunostaining quantification (n=4/group) (**C-D**). ADM: acinar to ductal metaplasia. (**E**) Schematic of spontaneous KPPC treatment experiments. Treatment was initiated at 6 weeks and mice euthanized at 9 weeks post-birth. (**F-G**) Representative H&E images of the pancreata of KPPC mice (**F**) and quantification (Treatment initiation: n=4, Vehicle^MRTX1133^: n=1 and MRTX1133: n=1) (**G**). In **C** and **D**, data are presented as mean + SD. Significance was determined by two-way ANOVA with Sidak’s multiple comparisons test in **C** and by unpaired-T test in **D**. Scale bar: 100 μm. ** P<0.01, **** P<0.0001.

Next, autochthonous genetic models of invasive PDAC with expression of *Kras^G12D^* and loss of *Trp53* (KPPC) and advanced disease were treated with MRTX1133 or vehicle control (**Fig. 3E**). Treatment for 3 weeks demonstrated a reversion of PDAC to predominantly uninvolved pancreatic tissue and PanIN histology (**Fig. 3F-G**), suggesting that inhibition of Kras* is sufficient to reprogram cancer cells to a premalignant state and potentially enable the regeneration of normal pancreatic parenchyma.

### Kras* targeting is associated with reprogramming of stromal fibroblasts and immune cells in the PDAC microenvironment

scRNA-seq analysis of KPC689 tumors in C57BL6/J mice was conducted to evaluate the impact of Kras* targeting with MRTX1133 on the tumor microenvironment (**Fig. 4A, Supplemental Fig. 4A-B**). scRNA-seq analysis revealed a proportional increase in fibroblasts upon treatment with MRTX1133 (**Fig. 4B**). MRTX1133 treated KPC689 tumors in C57BL6/J mice demonstrated increased podoplanin^+^ fibroblasts (**Fig. 4D**), a reported pan-fibroblast marker in PDAC (Elyada et al., 2019). Evaluation of stromal fibroblast clusters in KPC689 tumors implanted in NSG and C57BL6/J mice revealed transcriptionally distinct subsets of fibroblasts (**Fig. 4E-F, Supplemental Fig. 4C-E, 5A-D**) as previously reported (McAndrews et al., 2022). The diversity of CAFs subclusters were more pronounced in immune deficient (NSG) mice compared to immunocompetent mice with PDAC (C57BL/6J and KPPC mice), though in all groups MRTX1133 yield a shift in relative proportion toward aCAFs and bCAFs (McAndrews et al., 2022), which in part captures under a distinct nomenclature iCAFs (Elyada et al., 2019) (**Fig. 4E-H, Supplementary Fig. 5E**). Clustering of fibroblasts based on the previously reported myofibroblastic CAFs (myCAFs), inflammatory CAFs (iCAFs), and antigen presenting CAFs (apCAFs) (Elyada et al., 2019) revealed a shift to predominantly iCAFs with MRTX1133 treatment (**Supplemental Fig. 5E-H**). Notably, a/bCAFs and iCAFs are associated with early stage of disease (McAndrews et al., 2022). Consistently with this observation, stromal fibroblasts in C57BL6/J KPC689 and KPPC tumors treated with MRTX1133 displayed a proportional distribution similar to healthy pancreas, whereas fibroblasts in NSG KPC689 tumors acquired an intermediate state between fibroblasts of healthy and tumor bearing tissue (**Fig. 4H**), suggesting that CAFs accumulation and heterogeneity in PDAC progression and therapy response may be regulated by tumor infiltrating immune cells.

**Figure 4:**
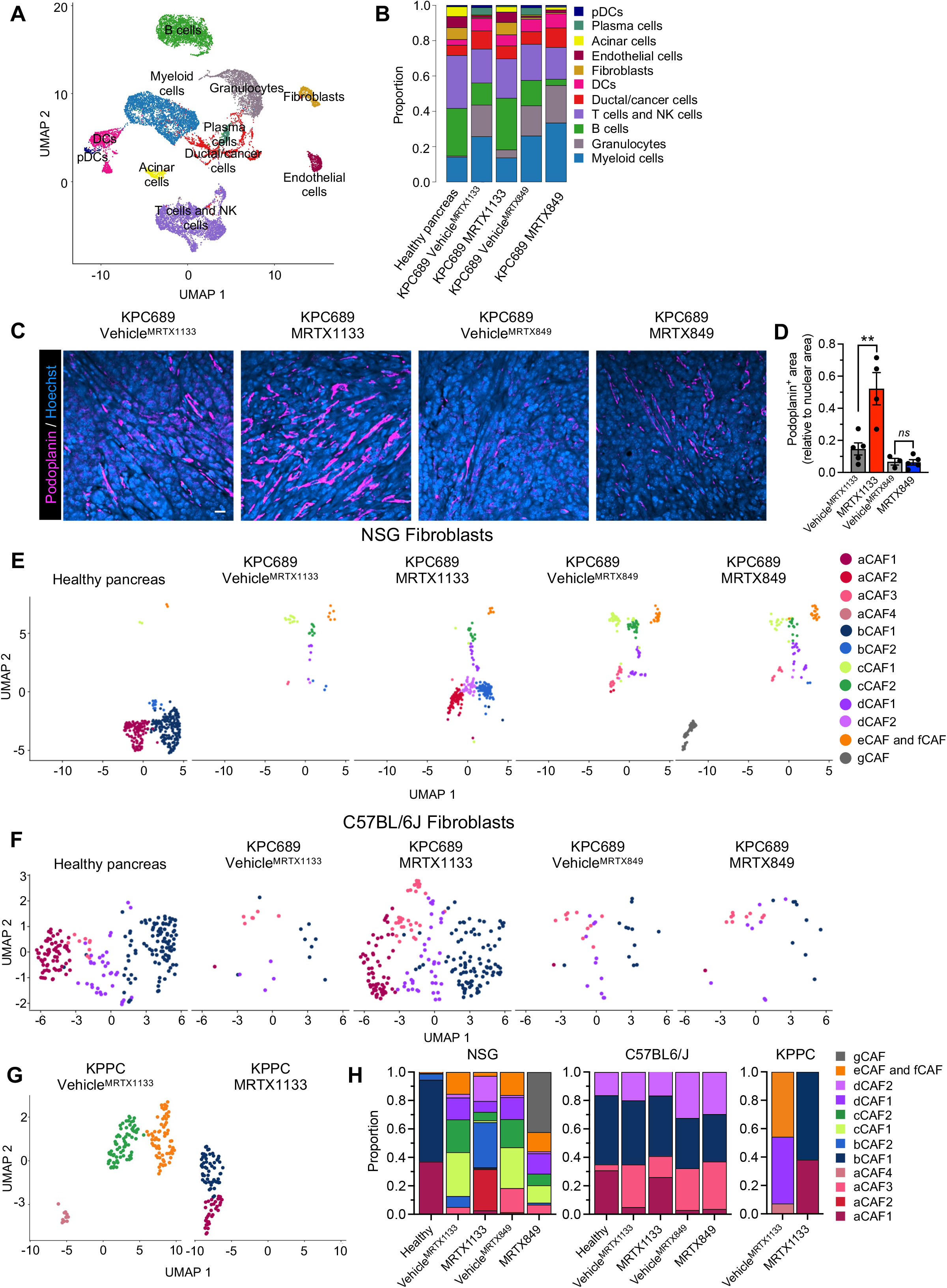
Fibroblast composition is altered in an immune-dependent manner in response to MRTX1133. (**A-B**) UMAP projection (**A**) and relative proportions of cell types (**B**) of orthotopic KPC689 tumors in C57BL6/J mice determined by scRNA-seq. (**C-D**) Representative immunofluorescent images of podoplanin stained orthotopic KPC689 tumors in C57BL6/J mice (**C**) and quantification of podoplanin^+^ area relative to nuclear area (**D**). Scale bar, 25 μm. (**E-G**) UMAP projections of fibroblasts in orthotopic KPC689 tumors in NSG mice (E) and C57BL6/J mice (**F**) and spontaneous KPPC tumors (**G**) determined by scRNA-seq. (**H**) Relative proportions of fibroblast subsets in orthotopic KPC689 tumors in NSG mice and C57BL6/J mice, and spontaneous KPPC tumors. Data are presented as mean in **B, H** and as mean + SD in **D**. Significance was determined by unpaired-T test in **D**. ** P<0.01, *ns*: not significant.

MRTX1133 treatment resulted in a significant increase in the frequencies of CD3^+^ T cells, CD8^+^ T cells, and CD19^+^ cells, as ascertained by immunotyping analyses (**Fig. 5A**). Analysis of T cell subsets by scRNA-seq revealed that MRTX1133 reduced the relative proportion of NK cells and increased the relative proportions of naïve CD4^+^,CD8^+^, and CD8^+^ effector T cells in both the KPPC mice and the orthotopic KPC689 model (**Fig. 5B-D, Supplemental Fig. 6A-B**). Further analysis of effector CD8^+^ T cell population revealed increased expression of *Ifng* (encoding IFNγ), *Prf1* (encoding perforin) and *Tbx21* (encoding T-bet) in the MRTX1133 treated tumors (**Fig. 5E, Supplemental Fig. 6C**), suggestive of accumulation of activated T cells. Immunostaining confirmed an increase in Ki67^+^CD8^+^ T cells upon MRTX1133 treatment (**Fig. 5F-G**). MRTX1133 reduced frequencies of CD11b^+^ myeloid infiltration as seen by flow cytometer, immunohistochemistry and sc-RNA seq analysis (**Fig. 5A, Supplemental Fig. 6D-F**). Further analysis of other myeloid cell subsets or macrophage infiltration showed that although MRTX1133 suppressed *Arg1^+^* and enriched *Mrc1^+^* macrophage clusters (seen in scRNA-seq analysis), such shifts did not impact the overall frequency of CD11b^+^F4/80^+^ macrophage infiltration (**Fig. 5A, Supplemental Fig. 6D-F**). Insignificant changes in CD11c^+^, CD11c^+^F4/80^-^, and CD3^-^NK1.1^+^ cell infiltration were observed upon treatment with MRTX1133 treatment (**Supplemental Fig. 6G-I**).

**Figure 5:**
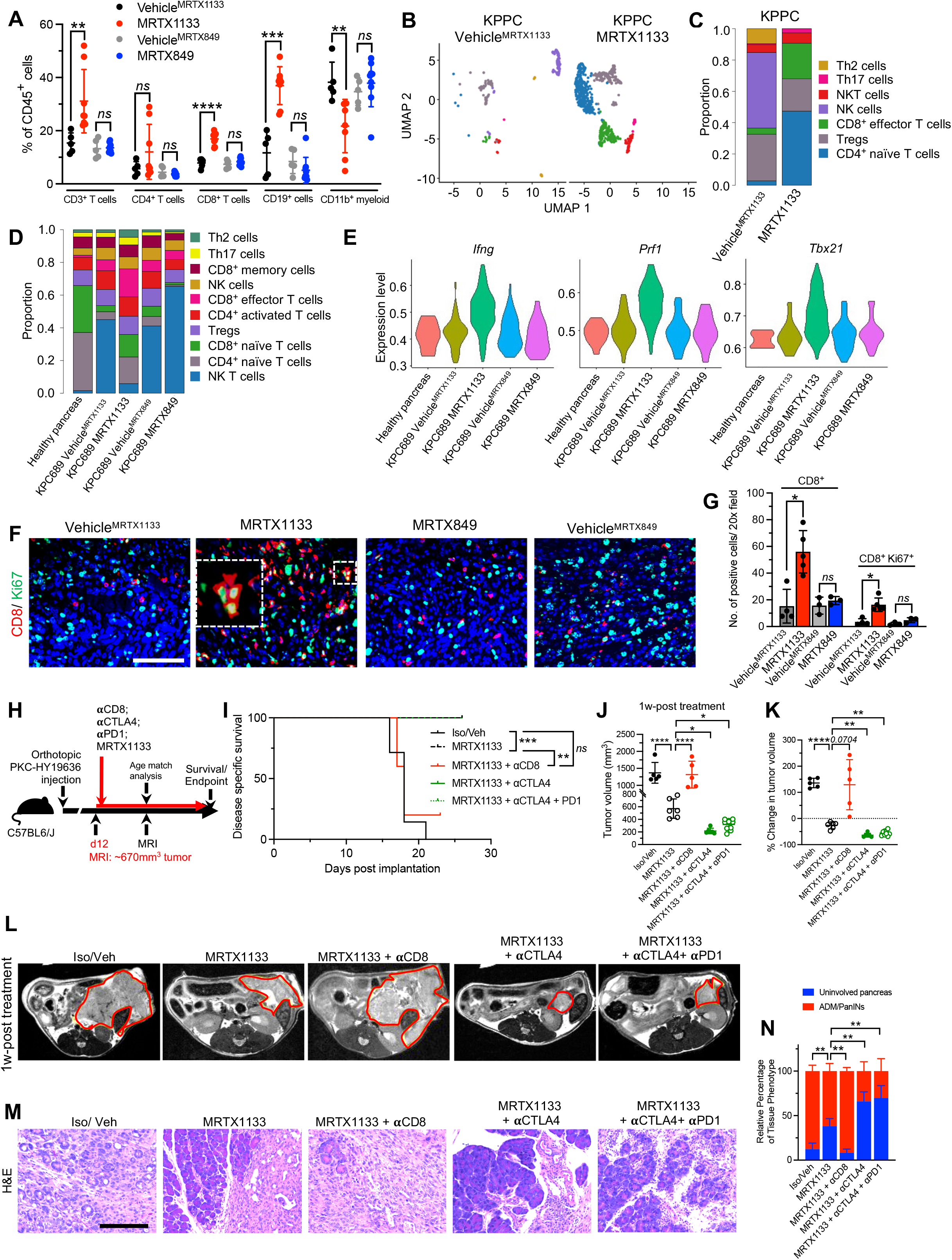
The efficacy of MRTX1133 is dependent on CD8^+^ T cells. (**A**) Immunotyping by flow cytometry of orthotopic KPC689 tumors in C57BL6/J mice (Vehicle^MRTX1133^: n=5, MRTX1133: n=8, Vehicle^MRTX849^: n=5, MRTX849: n=9). (**B-C**) UMAP projection and relative abundance of T cell subsets (**C**) of spontaneous KPPC tumors determined by scRNA-seq. (**D**) Relative abundance of T cell subsets of orthotopic KPC689 tumors in C57BL6/J mice determined by scRNA-seq. (**E**) Violin plots for *Ifng, Prf1,* and *Txb21* expression in tumor infiltrating CD8^+^ effector T cells in orthotopic KPC689 C57BL6/J mice determined by scRNA-seq. (**F-G**) Representative immunofluorescent images (**F**) and quantification (**G**) of CD8^+^ and Ki67^+^ T cells in orthotopic KPC689 tumors in C57BL6/J mice. (**H**) Schematic of experiments with C57BL6/J mice orthotopically implanted with PKC-HY19636. Treatment started when tumors were 670 mm^3^ by MRI. (**I**) Survival of mice with PKC-HY19636 orthotopic tumors with the indicated treatment groups (Iso/ veh (n=7), MRTX1133 (n=7), MRTX1133 + αCD8 (n=8), MRTX1133 + αCTLA4 (n=8), MRTX1133+ αCTLA4 + αPD1(n=10)) (**J**) Tumor volumes at 1 week post treatment initiation in the indicated treatment groups by MRI (Iso/ veh (n=5), MRTX1133 (n=6), MRTX1133 + αCD8 (n=5), MRTX1133 + αCTLA4 (n=6), MRTX1133+ αCTLA4 + αPD1(n=10)). (**K**) Change in tumor volumes of indicated groups compared to baseline by MRI (Iso/ veh (n=5), MRTX1133 (n=6), MRTX1133 + αCD8 (n=5), MRTX1133 + αCTLA4 (n=6), MRTX1133+ αCTLA4 + αPD1(n=10)). (**L**) Representative MRI of tumor burden 1 week post treatment initiation. (**M**) Representative H&E of the pancreas in the indicated group and (**N**) associated histopathological scoring (Iso/ veh (n=5), MRTX1133 (n=3), MRTX1133 + αCD8 (n=3), MRTX1133 + αCTLA4 (n=3), MRTX1133+ αCTLA4 + αPD1(n=3)). Data are presented as mean + SD in **A, G, J, K** and **N**; as mean in **C** and **D**, and as violin plots in **E**. In **A**, Mann-Whitney test was used for all comparison of CD4^+^ T cells and for MRTX1133 comparisons in CD3^+^ T cells and CD19^+^ cells. Unpaired-T test was used for all other comparisons in **A**. In **G**, Mann-Whitney test was used for MRTX1133 comparisons and unpaired-T test was used for all other comparisons. In **J** and **K**, one-way ANOVA with Holm-Sidak’s multiple comparisons test was used to determine significance. In **N**, two-way ANOVA with Sidak’s multiple comparisons test was used to determine significance. In **I**, log rank test was used to determine significance. * P<0.05, ** P<0.01, *** P<0.001, **** P<0.0001, *ns*: not significant. Scale bar: 100 μm.

To evaluate the functional contribution of immune cells and assess if tumor infiltrating T cells following MRTX1133 can render PDAC sensitive to iCBT, the PKC-HY19636 cell line was orthotopically implanted and treatment with MRTX1133, in combination with CD8 depletion, or iCBT (αCTLA4 and αPD1), was initiated in advanced PDAC stage, with a starting tumor volume of 670 mm^3^ captured by MRI (**Fig. 5H, Supplemental Fig. 7A-B**). After 1 week of treatment, all vehicle-treated control mice and MRTX1133 plus anti-CD8 treated mice succumbed to PDAC with an average tumor size of 1,350 mm^3^ (**Fig. 5I-L**). All MRTX1133 alone treated mice were still all alive after 1 week of treatment with average tumor size of 570 mm^3^. All MRTX1133 plus anti-CTLA-4 iCBT or plus anti CTLA-4+ anti-PD-1 iCBT treated mice were still all alive after 1 week of treatment with average tumor size of 275 mm^3^ (**Fig. 5J**). Serial MRI measurements show a regression of tumor volume when MRTX1133 is given in combination with iCBT compared to MRTX1133 as single agent or other controls (**Fig. 5K, L**). Further analysis of CD8^+^ T cells demonstrated systemic and intra-tumoral depletion in the mice treated with anti-CD8 antibody and MRTX1133 **(Supplemental Figure S7D-F)**. Although there was a modest increase in intra-tumoral CD8^+^ T cells in the MRTX1133 treated mice, addition of anti-CTLA4 and anti-CTLA4+ anti-PD1 iCBT demonstrated a robust influx of CD8^+^ T cell infiltrates and enhanced tumor regression in these mice compared to MRTX1133 monotherapy (**Fig. 5I, M, N and Supplemental Fig. 7E-F**). Depletion of CD8^+^ T cells with MRTX1133 abrogate the anti-tumor efficacy of the Kras* inhibitor (**Fig. 5K**). Taken together, the data support that eradication of established, large tumor with Kras* inhibition requires CD8^+^ T cells, and iCBT enables immune clearance of MRTX1133 treated tumors.

### Kras* inhibition upregulates Fas in cancer cells to facilitate CD8^+^ T cell mediated apoptosis

Efficacy of MRTX1133 in advanced PDAC is dependent on CD8^+^ T cell mediated clearance of cancer cells (**Fig. 5I**). To evaluate the mechanism(s) by which cancer cells become sensitive to adaptive immune response, scRNA-seq analysis of cultured KPC689 cells treated with MRTX1133 was performed. A shift in transcriptional clusters was observed with MRTX1133 treatment, with cluster 0 and 7 becoming the most prevalent clusters in MRTX1133 treated cells (**Fig. 6A-B**). Pathway analysis revealed that Kras signaling, Myc targets, and mTORC1 signaling were suppressed in these clusters (**Supplemental Fig. 7G-H**), supporting effective targeting of Kras* (Soucek et al., 2008; Ying et al., 2012). In addition, several immune modulatory pathways were suppressed in MRTX1133 treated clusters, including TNFα signaling via NFκB, inflammatory response, and allograft rejection (**Supplemental Fig. 7G-H**). Further analysis of death receptors in KPC689 cells revealed that *Fas* expression was increased upon MRTX1133 treatment (**Fig. 6C**). Moreover, MRTX1133 treated orthotopic KPC689 tumors revealed an increase relative proportions of CD8^+^ cells expressing *Fasl* (**Fig. 6D, Supplemental Fig. 7I**), supporting the possibility of CD8^+^ T cell mediated cancer cell clearance through Fas-FasL interaction. Treatment of PDAC cell lines with AZD6224 (a MEK inhibitor) also significantly increased *FAS* expression (**Fig. 6E-F**). MRTX1133 and MRTX849 induced Fas expression in Kras* and Kras^G12C^ mutant cell lines, respectively (**Fig. 6G-I**). Similar induction of *FAS* expression in response to MRTX849 and MRTX1133 was observed in a PDAC patient derived organoid (PDO, line DH-50) harboring both *KRAS^G12D^* and *KRAS^G12C^* mutations **Fig. 6J-O**).

**Figure 6:**
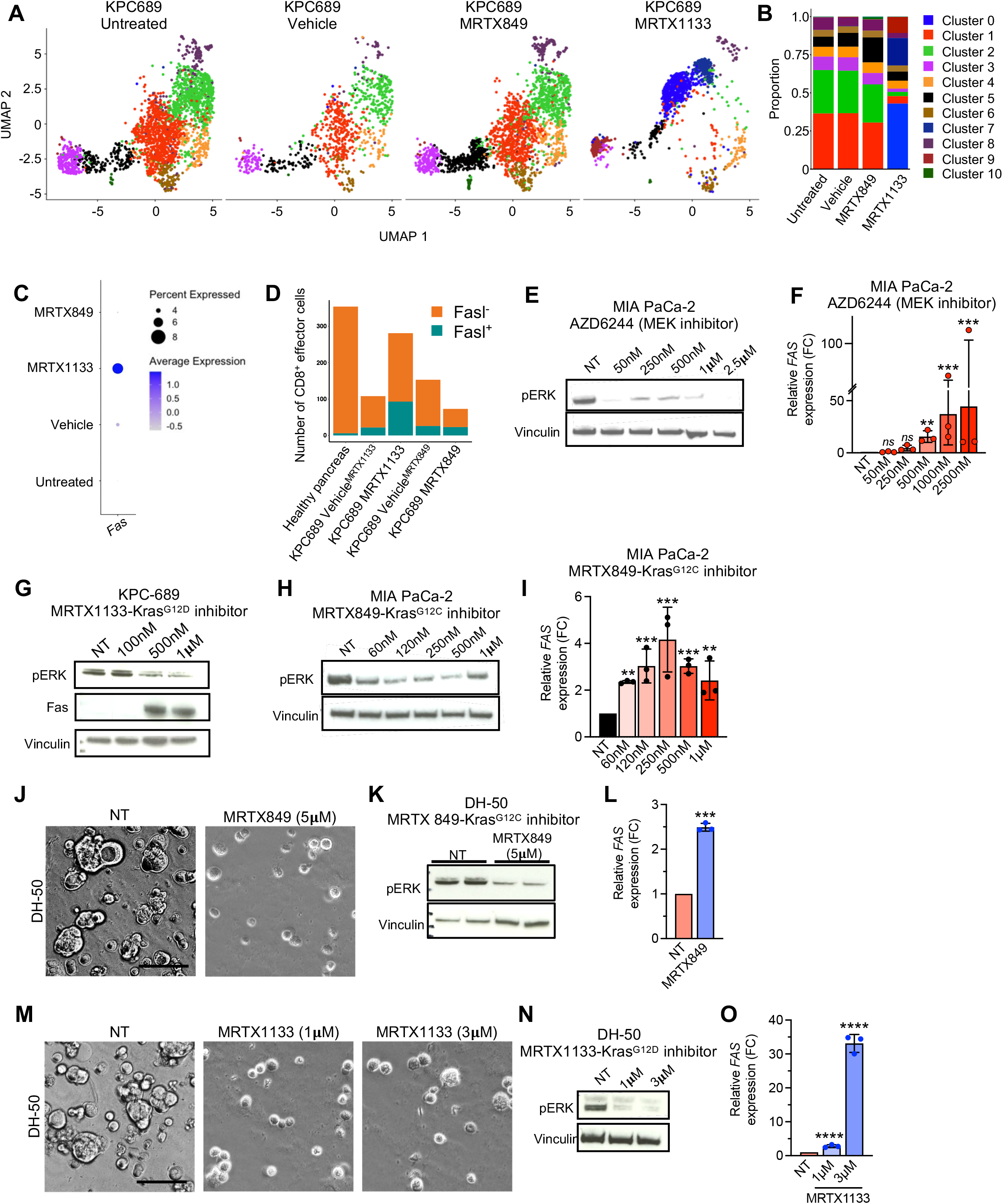
Mutant Kras inhibition induces Fas expression in cancer cells. (**A**) UMAP projections and relative proportions of each cluster (**B**) of KPC689 cells evaluated by scRNA-seq analysis. (**C**) Expression of *Fas* in KPC689 cells evaluated by scRNA-seq analysis. (**D**) Number of CD8^+^ effector cells with *Fasl* (Fasl^+^) or without *Fasl* (Fasl^-^) expression in KPC689 tumors in C57BL6/J mice evaluated by scRNA-seq analysis. (**E**) Western blot analysis of pERK expression level in MIA PaCa-2 cells treated with MEK inhibitor (AZD6244). (**F**) qPCR analysis of relative FAS expression in MIA PaCa-2 cells treated with MEK inhibitor (AZD6244) (n=3 biological replicates per group). (**G**) Western blot analysis of pERK and Fas expression level in KPC689 cells treated with Kras^G12D^ inhibitor (MRTX1133). (**H**) Western blot analysis of pERK expression level in MIA PaCa-2 cells treated with Kras^G12C^ inhibitor (MRTX849). (**I**) qPCR analysis of relative FAS expression in MIA PaCa-2 cells treated with Kras^G12D^ inhibitor (MRTX1133) (n=3 biological replicates per group). (**J**) DH-50 organoids were treated with Kras^G12C^ inhibitor (MRTX849) (n=3 biological replicates per group). Representative phasecontrast microscope image. (**K**) Western blot analysis of pERK expression in DH-50 organoids treated with Kras^G12C^ inhibitor (MTX849). (**L**) qPCR analysis of relative FAS expression in DH-50 organoids treated with Kras^G12C^ inhibitor (MRTX849) (n=3 biological replicates per group). (**M**) DH-50 organoids were treated with Kras^G12D^ inhibitor (MRTX1133) (n=3 biological replicates per group). Representative phase-contrast microscope image. (**N**) Western blot analysis of pERK expression in DH-50 organoids treated with Kras^G12D^ inhibitor (MTX1133). (**O**) qPCR analysis of relative FAS expression in DH-50 organoids treated with Kras^G12D^ inhibitor (MRTX1133) (n=3 biological replicates per group). Data are presented as mean + SD in **F, I, L** and **O,** and as mean in **B** and **D.** Significance was determined by one-way ANOVA with Dunnett’s multiple comparisons test in **F, I** and **O,** and by unpaired-T test in **L**. *P<0.05, **P<0.01, ***P<0.001, ****P<0.0001, *ns*: not significant.

Suppression of the *Kras\** transcript with siRNA (siKras^G12D^) in *Kras^G12D^* mutant PDAC cells was also associated with increased *FAS* expression, likely through reversal of Kras* mediated transcriptional repression of Fas by enrichment of DNMT1, EZH2, and H3K27me3 and increased enrichment of H3K27Ac on the Fas promoter (**Fig. 7A-B**). To define whether targeting of *Kras*\* mRNA could induce Fas-Fasl mediated apoptosis, as observed with pharmacological inhibition with MRTX1133, orthotopic KPC689 tumors and autochthonous *P48-Cre; LSL-Kras^G12D/+^; Tgfbr2^L/L^* (PKT) mice were treated with exosomes containing Kras^G12D^ siRNA (iExo^siKrasG12D^) (Kamerkar et al., 2017; Mendt et al., 2018). iExo^siKrasG12D^ was associated with increased CD4^+^ and CD8^+^ T cell infiltration (**Fig. 7C-D**) and increased the number of Fas^+^CK19^+^ cancer cells (**Fig. 7E-F**). The impact of CD8^+^ T cells on the efficacy of iExo^siKrasG12D^ was evaluated in the context of *LSL-Kras^G12D/+^; LSL-Trp53^R172H/+^; Pdx1-Cre* (KPC) CD8^-/-^ mice which lack CD8^+^ T cells and develop spontaneous PDAC. Mice were enrolled for iExo^siKrasG12D^ treatment at 14 weeks of age, when tumors were detectable by MRI (**Fig. 7G-H**). KPC mice lacking CD8^+^ T cells that were treated with iExo^siKrasG12D^ displayed significantly reduced survival and increased tumor volume (**Fig. 7G-I**), similar to the MRTX1133 treated mice (**Fig. 5I**). Similarly, survival was decreased in CD8^-/-^ mice implanted with orthotopic KPC689 tumors and treated with iExo^siKrasG12D^ (**Fig. 7J**), as also observed in the context of Kras* targeting with MRTX1133 (**Fig. 6I**). Together these data indicate that clearance of cancer cells following pharmacological inhibition or genetic inhibition of Kras^G12D^ requires CD8^+^ T cell mediated killing.

**Figure 7:**
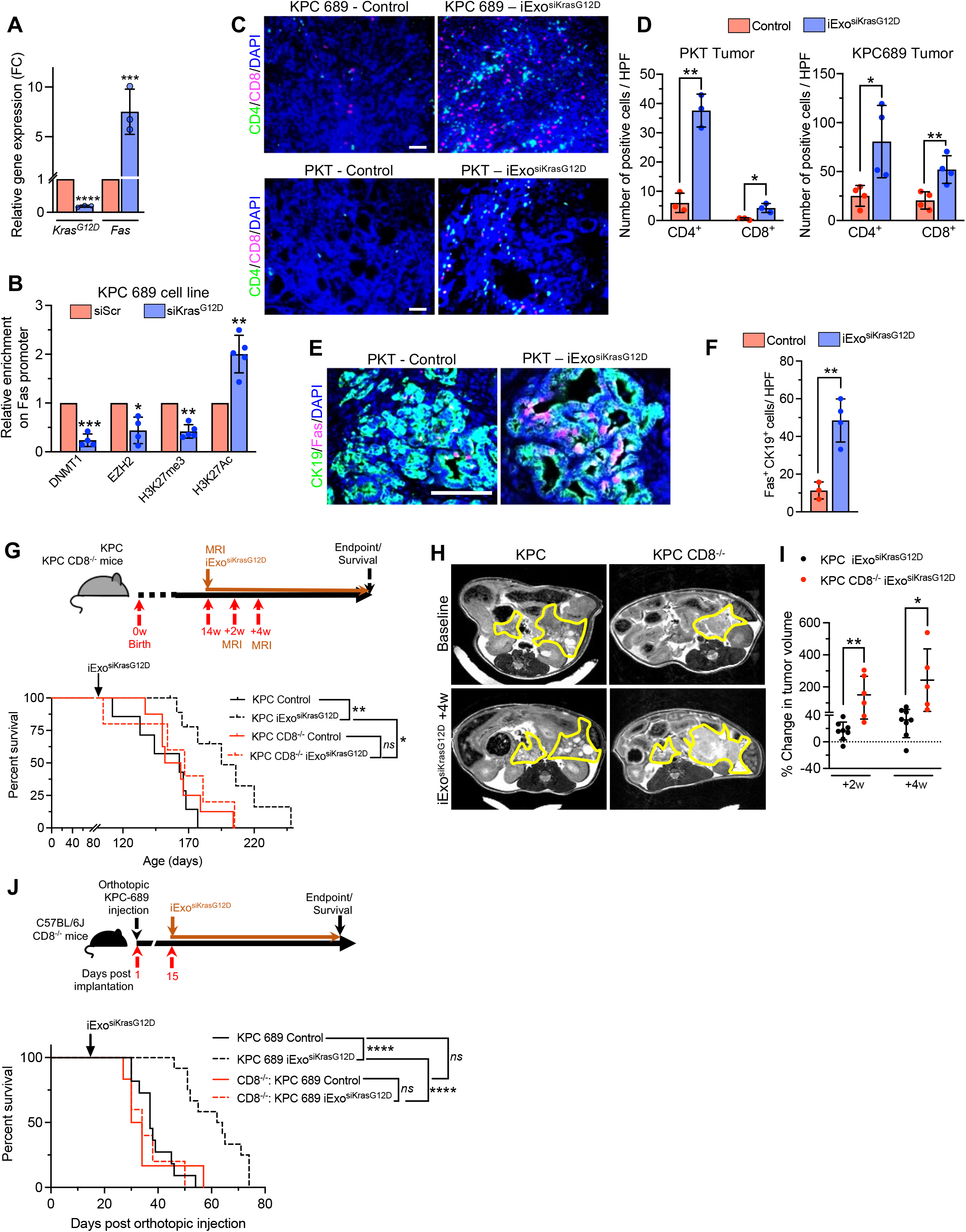
CD8^+^ T cells mediate the efficacy of iExo^siKrasG12D^. (**A**) qPCR analysis of relative *Kras^G12D^* and *Fas* expression in KPC-689 cells 72 hours after siKras^G12D^/ siScr treatment (n=3 biological replicates per group). (**B**) ChIP assays demonstrating the relative binding of DNMT1, EZH2, H3K27me3 and H3K27ac on Fas promoter in KPC-689 cell line following siKrasG12D treatment (n=4 biological replicates/ group for DNMT1, EZH2 ChIPs and n=5 biological replicates/ groups for histone ChIPs). (**C-D**) CD4 and CD8 immunolabeling – control (n=4), KPC 689 – iExo (n=4) treated tumors (top panel) and age matched (44 days) PKT (n=3) and PKT iExo (n=3) treated tumors (bottom panel). (**E-F**) CK19 and Fas immunolabeling (E) and quantification (F) of age matched (44 days) PKT (n=3) and PKT iExo (n=4) treated tumors. (**G**) (Top panel) Timeline of the experiment for KPC and KPC CD8^-/-^ mice treated with iExosomes (iExo) every 48h starting at 14 weeks of age. Baseline MRI measurements were done prior to enrolment into iExo treatment groups and were subsequently imaged at indicated time points. (Bottom panel) Kaplan Meier survival curves of KPC Control (n=7), KPC iExo (n=11), KPC CD8^-/-^ Control (n=8), KPC CD8^-/-^ iExo (n=5) mice. (**H**) Baseline (top) and 4 weeks post iExo time point (bottom) MRI imaging of tumors following iExosomes treatment. (**I**) Change in tumor volumes (based on MRI measurements) of KPC and KPC CD8^-/-^ at 2 weeks and 4 weeks following iExo treatment (KPC iExo +2w (n=8), KPC iExo +4w (n=8), KPC CD8^-/-^ iExo +2w (n=6), KPC CD8^-/-^ iExo +4w (n=5)). **(J)** (Top panel) Timeline of the experiment. WT-C57BL6/J and CD8^-/-^ mice were injected with KPC-689 cells. iExo treatment was started in both the cohorts on day 14. (Bottom panel) Kaplan-Meier survival curves of KPC 689 Control (n=11), KPC 689 iExo (n=12), CD8^-/-^: KPC 689 Control (n=6), CD8^-/-^: KPC 689 iExo (n=5) mice. In all images, scale bars, 100 μm. In **(B, C, D, F** and **I)** data are presented as the mean ± SD. Significance was determined by unpaired T-test in **(B, C, D, F** and **I),** log rank test in **(G** and **J)**. * P<0.05, ** P<0.01, ***P<0.001, *ns*: not significant.

## Discussion

Recent identification of a non-covalent small molecule inhibitor (MRTX1133) with specificity to the Kras^G12D^ mutant protein has offered an opportunity to evaluate its efficacy and mechanism of action directly on Kras* cancer cells and the entire PDAC tumour (Hallin et al., 2022; Kemp et al., 2023). Here, we study MRTX1133 in 15 different mouse models of pancreatic cancer, which include orthotopic xenografts, orthotopic syngeneic models, patient derived xenografts, KC genetic model, and KPPC genetic model. The goal was to focus on the efficacy of MRTX1133 against early and late stage PDAC and determine the impact of this inhibitor on Kras* PDAC cells, along with its impact on the tumor microenvironment, with a specific focus on immune cells and stromal fibroblasts.

We show that MRTX1133 has significant impact on the growth of early stage tumors with tumor stasis observed in all human xenograft models, as also shown in other recent studies(Kemp et al., 2023). We demonstrate that MRTX1133 inhibits proliferation of PDAC cells and suppresses pERK, and such activity has a dominant impact on inducing tumor inhibition; however, for sustained suppression and tumor clearance, CD8^+^ T cells are required. The critical function of CD8^+^ T cells in tumor clearance became apparent when treatment with MRTX1133 was initiated in mice with large tumor burden associated with advanced PDAC. In this regard, we show that genetic extinction of Kras* (Mahadevan et al 2023 in review) or suppression of Kras* with MRTX1133 or iExo^siKrasG12D^ results in upregulation of Fas via demethylation of Fas promoter on PDAC cells, rendering them vulnerable to FasL mediated apoptosis by CD8^+^ T cells (Mahadevan et al 2023 in review).

While other studies have suggested that CD8^+^ T cells are not required for suppression of tumor growth in subcutaneous and orthotopic PDAC tumors, they involve short term experiments using mice with small tumor burden(Kemp et al., 2023). In this setting, it is likely tumor stasis due to suspended proliferation can appear effective but if all the cancer cells are not eliminated, tumor reemergence is possible. Our study demonstrates that MRTX1133 is effective in regression of established tumors associated with advanced PDAC. Such activity depends on the activity of CD8^+^ cells and their suppression leads to rapid tumor growth despite treatment with MRTX1133. Further, Kras* inhibition with iExo^siKrasG12D^ highlights the critical role of CD8^+^ T cells in suppression of tumor growth following Kras* targeting therpy. These experiments suggest that evaluation of the efficacy of MRTX1133 in pre-clinical studies must involve mouse models with different tumor growth kinetics that include both early and late stage PDAC. Using this rationale, our study uniquely identifies a dominant impact of MRTX1133 in regressing pre-established, advanced and aggressive PDAC with large tumor burden. Guided by the data that demonstrates the emergence of T cells upon treatment with MRTX1133, we evaluated the combination of MRTX1133 with anti-CTLA4 and anti-CTLA-4+anti-PD1 iCBT. These results show that a combination of MRTX1133 with iCBT is more efficient in tumor regression when compared to MRTX1133 alone.

Collectively, our studies demonstrate that MRTX1133 is specific and efficient inhibitor of Kras* and with the ability to regress PanINs and advanced PDAC in autochthonous and orthotopic models of PDAC. MRTX1133 treatment reprograms tumor immunity and stromal fibroblasts and synergizes with iCBT to further regress tumors with increase in overall survival of treated mice. Such combination treatment opportunity informs current clinical development of MRTX1133.

## Methods

### Kras^G12D^, Kras^G12C^ and MEK small molecule inhibitors

MRTX849 (adagrasib) was purchased from MedChem Express and MEK inhibitor AZD6224 was purchased from Stem Cell Technologies. MRTX1133 was synthesized by WuXi AppTech. MRTX849 and MRTX1133 compounds were structurally verified by NMR and LC/MS. For cell culture experiments, MRTX849, MRTX1133, and AZD6244 were diluted in dimethyl sulfoxide (DMSO).

### Cell culture

Cell lines employed in these studies, their origin, and culture conditions are listed in **Table 1**. KPC689 cells were isolated from a spontaneous *LSL-Kras^G12D/+^* (Jackson *et al.,* 2001); *LSL-Trp53^R172H/+^* (Olive et al., 2004); *Pdx1-Cre* (Hingorani et al., 2003) (KPC) tumor as previously described (Kamerkar et al., 2017; Zheng et al., 2015), serially passaged in C57BL6/J mice. PKC-HY19636 cells were isolated from a spontaneous *P48-Cre; LSL-Kras^G12D/+^, Trp53^L/+^* tumor as previously described (Deng et al., 2021). Panc1 and KPC689 were engineered to stably express GFP and firefly luciferase by lentiviral infection (Capital Biosciences) and maintained in 2 μg/mL puromycin (Sigma Aldrich). Cells were routinely tested for mycoplasma with LookOut Mycoplasma PCR Detection Kit (Sigma Aldrich) and confirmed to be mycoplasma negative. HPAC, Panc1, A549, HCT116, MIA PaCa-2, and PSN-1 were STR verified by the MDACC Cytogenetics and Cell Authentication Core.

**Table 1:** Cell lines and culture conditions.

| Cell line | Origin | Media |
| --- | --- | --- |
| HPAC | ATCC | DMEM/F12 (Corning), 5% FBS (Gemini), 0.002 mg/mL insulin (Sigma Aldrich), 0.005 mg/mL transferrin (Sigma Aldrich), 40 ng/mL hydrocortisone (Sigma Aldrich), 10 ng/mL EGF (Peprotech), 1% penicillin-streptomycin (P/S) |
| Panc1 | ATCC | RPMI (Corning), 10% FBS, 1% P/S |
| A549 | ATCC | DMEM (Corning), 10% FBS, 1% P/S |
| HCT116 | ATCC | McCoy's-5A (Corning), 10% FBS, 100 mM L-glutamine, 1% NEAA, 1% P/S |
| PSN-1 | ATCC | RPMI (Corning), 10% FBS, 1% P/S |
| MIA<br>PaCa-2 | ATCC | RPMI (Corning), 10% FBS, 1% P/S |
| KPC689 | (Kamerkar et al., 2017; Zheng et al., 2015) | RPMI (Corning), 10% FBS, 1% P/S |
| PKC-HY19636 | (Deng et al., 2021) | DMEM (Corning), 10% FBS, 1% P/S |

### Patient derived organoids

The DH-50 PDO was generated from a liver metastasis lesion of a PDAC patient and expanded as a patient derived xenograft. PDO cultures and expansion was performed as described previously (Huang et al., 2015). Briefly, PDO was cultured in Nunclon™ Delta Surface (Thermoscientific) on Geltrex dome (Reduced growth factor basement membrane, hESV qualified, Gibco) suspended in PaTOM growth media (Huang et al., 2015). TrypLE Express (Gibco) was used for passage of organoids and DMEM, 10% FBS and 1% P/S media was used to neutralize TrypLE.

### Cell proliferation assays

Cells were plated at the density of approximately 2,500 cells/well in clear-bottom, white-wall 96-well plates (Corning) in DMEM (Gibco) supplemented with sodium pyruvate (Sigma Aldrich), PS and 2.5% FBS. After 24 hours, old media was removed and replaced with a serial dilution of MRTX1133 starting from 10 μM for three days. The CellTiter-Glo (Promega) assay was used to assess the viability of cells.□CellTiter-Glo□reagents and plates were allowed to□equilibrate at room temperature, and 50 μl of reagent was added to 50□μl of media per well. The luminescent signal was read using a FLUOstar Omega plate reader (BMG LABTECH). Data were analyzed by normalizing the cell densities to the control wells. Constraints for IC50 calculations were set for the bottom to be less than 0.5 and top less than 1.0. Cell lines that did not have a calculable IC50 in the range of MRTX1133 evaluated are listed as ‘n.d.’ or not determined.

### Animal studies

All procedures were reviewed and approved by the Institutional Animal Care and Use Committee (IACUC) at MD Anderson Cancer Center (MDACC). Male and female, 6 to 16 weeks old C57BL/6J (KPC689 GFP Luc and PKC-HY19636 cells) for and NSG (for KPC689 GFP Luc, Panc-1 GFP Luc, or HPAC cells) mice were purchased from Jackson Laboratories and implanted orthotopically with KPC689 GFP Luc (0.5×10^6^ cells in 20 μL of PBS), PKC-HY19636 (0.5×10^6^ cells in 20 μL of PBS), Panc-1 GFP Luc, or HPAC, and (1×10^6^ cells in 20 μL of PBS) by injection into the tail of the pancreas using a 27-gauge Hamilton syringe. For detection of tumor burden, luciferase expression was measured by injecting the mice i.p. with D-Luciferin (Goldbio) per the manufacturer’s instructions and imaged via IVIS (Xenogen Spectrum). Alternatively, MRI imaging was performed on day 12 and followed up with serial measurements to assess tumor volumes using 7T Bruker MRI at the MDACC Small Animal Imaging Facility (SAIF). Mice were randomly assigned to groups and treatment was initiated at the indicated days post-surgery. Genetically engineered mice includiing *LSL-Kras^G12D/+^; Pdx1-Cre* (KC) and *LSL-Kras^G12D/+^; Trp53^L/L^* (Chen et al., 2005); *Pdx1-Cre* (KPPC) mice were maintained on mixed backgrounds and started on treatment at the indicated times post-birth. PDXs (**Table 2**) were derived as previously described (Carugo et al., 2016) and implanted subcutaneously into 11 weeks old athymic nude mice. Tumor volumes were captured by serial caliper measurements (twice weekly) and treatment was initiated when tumors reached 150 to 250 mm^3^. Tumor volume (TV) was calculated as TV = (D × d2/2), where “D” is the larger and “d” is the smaller superficial visible diameter of the tumor mass. All measurements were documented as mm^3^. Treatment groups included 6 mice per group. For acute pharmacodynamic (PD) biomarker studies (n=4 mice per group), tumors were allowed to grow to an average volume of 250–350 mm^3^ and collected 4 hours after the last dose of treatment and fixed in 10% neutral buffered formalin overnight and then processed and embedded in paraffin.

**Table 2:**
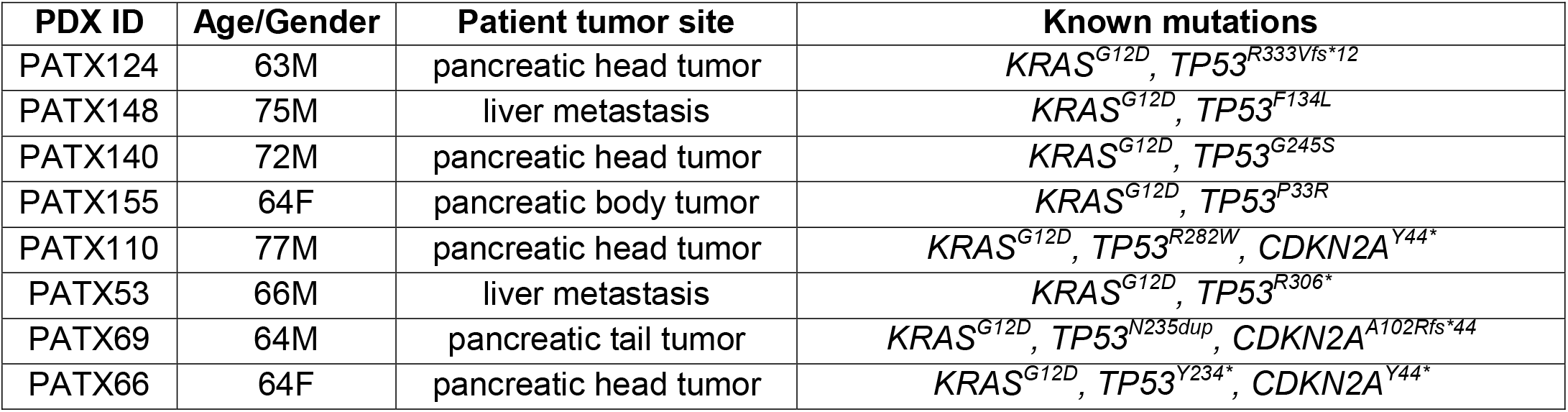
Patient derived xenografts (PDX).

| PDX ID | Age/Gender | Patient tumor site | Known mutations |
| --- | --- | --- | --- |
| PATX124 | 63M | pancreatic head tumor | *KRAS*<sup>G12D</sup>, *TP53*<sup>R333Vfs*12</sup> |
| PATX148 | 75M | liver metastasis | *KRAS*<sup>G12D</sup>, *TP53*<sup>F134L</sup> |
| PATX140 | 72M | pancreatic head tumor | *KRAS*<sup>G12D</sup>, *TP53*<sup>G245S</sup> |
| PATX155 | 64F | pancreatic body tumor | *KRAS*<sup>G12D</sup>, *TP53*<sup>P33R</sup> |
| PATX110 | 77M | pancreatic head tumor | *KRAS*<sup>G12D</sup>, *TP53*<sup>R282W</sup>, *CDKN2A*<sup>Y44*</sup> |
| PATX53 | 66M | liver metastasis | *KRAS*<sup>G12D</sup>, *TP53*<sup>R306*</sup> |
| PATX69 | 64M | pancreatic tail tumor | *KRAS*<sup>G12D</sup>, *TP53*<sup>N235dup</sup>, *CDKN2A*<sup>A102Rfs*44</sup> |
| PATX66 | 64F | pancreatic head tumor | *KRAS*<sup>G12D</sup>, *TP53*<sup>Y234*</sup>, *CDKN2A*<sup>Y44*</sup> |

Mice in control groups were administered 30 mg/kg MRTX1133 BID i.p. in 100 μL of 10% Captisol (Selleck) in 50mM sodium citrate pH 5 (Teknova, vehicle) or 30 mg/kg MRTX849 QD p.o. in 100 μL of vehicle. Vehicle^MRTX1133^ consisted of vehicle administered BID i.p. and Vehicle^MRTX849^ consisted of vehicle administered QD p.o. Concomitant with MRTX1133 (30 mg/kg, BID 5 days per week i.p.) treatment from day 13, depletion of CD8^+^cells was performed using 200 μg of anti-mouse CD8 (BioXcell, clone 53-6.7) twice per week (diluted to a final volume of 100 μL in PBS). For checkpoint immunotherapy, anti-mouse CTLA4 (BioXcell, clone 9H10) and/or anti-mouse PD-1 (BioXcell, clone 29F.1A12) were injected three times per week i.p., starting dose 200 μg followed by 2 doses of 100 μg each (diluted to a final volume of 100 μL in PBS). The control (Isotype/vehicle) mice received a cocktail of Rat IgG2a (BioXcell, clone 2A3), Syrian hamster IgG (BioXcell, BE0087) (200 μg of each antibody diluted in a final volume of 100 μL in PBS) and 100 μL of vehicle i.p.. The mice were euthanized at the indicated times post treatment.

Age matched PDAC tissue from iExosomes treated and control PKT and orthotopic KPC-689 tumor bearing mice were used for immunostaining analysis from a previous published study (Kamerkar et al., 2017). The generation and characterization of iExosomes was previously described (Kamerkar et al., 2017; Mendt et al., 2018). To generate KPC CD8^-/-^ mice, *Cd8a^tm1Mak^/J* (Fung-Leung *et al.,* 1991) (purchased from Jackson Laboratory) were crossed to KPC mice. Tumor volumes in the KPC and KPC CD8^-/-^ mice were assessed by Bruker 7T MRI as described above. For orthotopic KPC689 tumors, C57BL6/J or CD8^-/-^ mice (bred in C57BL6/J background) were injected with 1 x 10^6^ KPC689 cells in 20 μL PBS into the tail of the pancreas under general anesthesia. iExosomes treatment was started at 14 weeks in KPC, KPC CD8^-/-^ mice and at 2 weeks following orthotopic injection of KPC689 cells in C57BL6/J or CD8^-/-^ mice as described previously (Mendt et al., 2018). Briefly, mice were treated with 10^9^ GMP-compliant iExosomes (mesenchymal stromal cell derived exosomes electroporated with 1 μg siKras^G12D^-Avecia in plasmalyte) three times per week until mice reached endpoint. Non-electroporated, GMP-compliant exosomes were used as control exosomes.

### Histological analyses

3 to 5 μm thick sections were made from formalin fixed paraffin embedded (FFPE) blocks and were deparaffinized and hydrated. Antigen retrieval was performed in Tris-EDTA buffer (pH 9.0) for 20 minutes at 95 C in a steamer. Subsequently, a hydrophobic barrier was created around the tissue sections using Pap pen, and sections were incubated with 3% H_2_O_2_ in PBS for 15 minutes. For blocking, the slides were incubated in 1.5% bovine serum albumin in PBST (0.1% Tween 20) for 30 minutes. The sections were incubated with primary antibodies, CK19 (Abcam ab52625, 1:250), CD19 (Cell Signaling 3574s; 1:100), CD8 (Cell Signaling 85336S, 1:300), CD11b (Abcam ab133357, 1:1000), pERK (Cell Signaling 4376S, 1:250), diluted in blocking buffer overnight at 4 °C or for 3 hours at room temperature. For CD11b, pERK and CD8 immunohistochemistry, the slides were incubated in biotinylated anti-rabbit secondary (Vector Laboratories BA-1000, 1:250) for 30 minutes in blocking buffer followed by incubation in ABC reagent (Vector Laboratories PK-6100, prepared as per manufacturer’s instructions) for 30 minutes. Subsequently, slides were stained with DAB (Life Technologies), counterstained with haematoxylin, cover slipped and imaged with a Leica DM1000 LED microscope mounted with a DFC295 microscope camera (Leica) with LAS version 4.4 software (Leica). Multiple representative images were taken from each tissue section and quantified by counting the number of positive cells in each visual field or using Image J (NIH) for quantification of area.

For both acute PD studies and efficacy studies on PDAC PDX models, sections were subjected to an initial heat-induced epitope retrieval (HIER) in citrate buffer, pH 6, at 95°C for 15 minutes. Anti-phospho-p44/42 MAPK (ERK1/2) (Thr202/Tyr204) (1:2000, Cell Signaling Technology #4370) was developed using Opal tyramide signal amplification (TSA) followed by direct immunofluorescence of HLA conjugated to Alexa 647 (1:250, Abcam #199837). RNAscope in situ hybridization assay was performed following manufacturer’s protocol (Advanced Cell Diagnostics, Inc) using DUSP6 (cat# 405361), SPRY4 (cat# 546711-C2), and POLR2A (cat# 310451-C4) probes. Appropriate positive and negative controls were included with the study sections. Digital images of whole-tissue sections were acquired using Vectra Polaris Automated Quantitative Pathology Imaging System (Akoya Biosciences) and representative regions were selected for each whole slide and processed using inForm Software v2.4 (Akoya Biosciences). Processed images were then analyzed using HALO Software v3.2 (Indica Labs Inc.)

For CK19 and CD8 immunofluorescence staining, the slides were then incubated in rabbit-on rodent polymer (Biocare Medical) for 15 minutes following the primary antibodies. Subsequently, the sections were incubated in Opal 520 diluted in blocking buffer (for CK19 and CD8 primary) (Akoya Biosciences, 1:100) for 15 minutes. For CD8/Ki67 co-staining, another round of antigen retrieval was performed as described above and the sections were incubated in blocking buffer for 30 minutes and in Ki67 primary antibody (Thermo Fischer Scientific RM-9106-S, 1:100 concentration) diluted in blocking buffer overnight at 4 C. The slides were then incubated in rabbit-on rodent polymer for 15 minutes, followed by incubation in Opal 690 reagent for 15 minutes (Akoya Biosciences, 1:100) in blocking buffer. For CD4/CD8 co-staining, sections were incubated in CD4 primary antibody (Abcam ab183685, 1:300) following antigen retrieval and stained with rabbit-on rodent polymer and Opal 690 reagent as described above. For CK19/Fas co-staining, sections were incubated in Fas primary antibody (Abcam ab82419, 1:100) following antigen retrieval and stained with rabbit-on rodent polymer and Opal 690 reagent. Following another round of antigen retrieval, the slides were coverslipped with Fluoroshield with DAPI (Sigma Aldrich). PBST was used for washes between steps (3 mins each wash for 3 times). Imaging was performed with a Zeiss LSM800 confocal microscope with a 20x objective and quantified by counting the number of positive cells in each visual field.

For podoplanin immunofluorescence staining, antigen retrieval in TE buffer (pH 9.0) was performed for 15 minutes, followed by blocking in 1% BSA in TBS for 15 minutes. Slides were incubated in podoplanin primary antibody (Abcam ab1993, 1:200) for 1 hour at room temperature, washed with TBS for 2 minutes 3 times, then incubated with HRP goat anti-Syrian hamster IgG (Jackson Immunoresearch Laboratories 107-036-142, 1:250), washed with TBS for 2 minutes 3 times, and incubated in Opal 690 for 10 minutes. Antigen retrieval was repeated, followed by incubation in Hoechst 33342 (Invitrogen, 1:10,000) for 10 minutes and mounting with Vectashield mounting media (Vector Labs). Imaging was performed with a Zeiss LSM800 confocal microscope with a 20x objective and quantification of positive area performed in ImageJ (NIH).

For histopathological analysis of H&E stained tissues, pancreas/tumors were analysed under bright field microscopy with multiple low power (10x magnification) representative images to define the relative percentages of uninvolved-normal looking pancreas, acinar-ductal metaplasia (ADM) or Pancreatic intra-epithelial neoplasia (PanIN) leisons, and invasive PDAC lesions (classified further into well, moderate and poorly differentiated lesions). Liver necrosis was defined as areas of approximately than 5-10 hepatocytes or greater, and spot necrosis was defined as areas of 5 or less hepatocytes. Scores were set as 0 for no necrosis, 1 for 1-2 areas of necrosis detected, and 2 for 3 or more areas of necrosis detected per liver section. For binucleated cells, 5 periportal 400X field of view per mouse were selected at random and composite socres were defined as 0 for 1-2 binucleated cells, 1 for 3-5 binucleated cells, and 2 for greater than 6 binucleated cells per field of view. Kidney tubular damage was ascertained on 8 to 10 random 400X cortical fields of view and scored as 0 for no damaged tubules noted, 1 for 1 damaged tubule, and 2 for 2 or more damaged tubules. Kidney immune infiltration was ascertained on 5 to 8 random 400X cortical and papillary fields of view and scored as 0 for no significant (above healthy control) immune infiltrate, 1 for one or more fields of view with peritubular inflammation greater than observed in control (10-20 immune cells).

### Blood chemistry and urine analyses

Blood was collected without anti-coagulant via retro-orbital bleeding or cardiac puncture and incubated on ice for 20 minutes. The samples were then centrifuged at 6,000 rpm for 5 minutes at 4°C and the serum samples were submitted to the MDACC DVMS Veterinary Pathology Services for assessment of total bilirubin, albumin, alkaline phosphatase (ALP), alanine aminotransferase (ALT), aspartate aminotransferase (AST), blood-urea-nitrogen (BUN), lactate dehydrogenase (LDH), calcium, chloride, globulin, creatinine, potassium, sodium, phosphorus, glucose and total protein. Urine was collected and protein levels measured using URiSCAN test strips (BioSys Laboratories).

### Pancreas digestion and scRNA-seq

Pancreata/tumors were collected and combined from 3 mice for vehicle groups and MRTX849 treated mice and 6 mice for MRTX1133 treated mice. Pancreata/tumors were minced and digested at 37°C for 30 minutes at 150 rpm in a 10 mL mixture of 4 mg/mL collagenase IV (Gibco) and 4 mg/mL dispase (Gibco). Cells were filtered once with a 70 μm filter and once more with a 40 μm filter. To stop the digestion, 25 mL of DMEM containing 10% FBS was added. Cells were then centrifuged at 500 *g* for 5 minutes, and the supernatant was removed, washed with 10 mL of FACS buffer (2% FBS in PBS), and centrifuged at 500 *g* for 5 minutes at 4°C. Cells were resuspended in 1 mL of FACS buffer and stained with 1 μL of Fixable Viability Dye eFluor™ 780 (eBioscience, 1:1,000) on ice for 10 minutes. Cells were washed with 3.5 mL of FACS buffer, centrifuged at 300 g for 5 minutes at 4°C, washed again, and resuspended in 1 mL of FACS buffer. Live cells were subsequently sorted via the BD FACSMelody cell sorter, followed by counting with Trypan Blue (Sigma Aldrich) exclusion with a Countess 3 (Invitrogen). For in vitro scRNA-seq, cells were treated with vehicle (DMSO), 50 nM MRTX1133, or 50 nM MRTX849 for 3 hours then trypsinized, washed, and counted. 5,000 cells were loaded into the 10x Chromium Controller to generate Single Cell 3’ Gene Expression dual indexed libraries (10x Genomics). The creation of Gel Bead-In-Emulsions (GEM) and subsequent cell barcoding, GEM-RT cleanup, cDNA amplification, and library assembly were performed with the Chromium Next GEM Single Cell 3’ Reagents Kits v3.1 (10x Genomics) according to manufacturer’s instructions. Paired-end, dual indexing sequencing was carried out using Illumina High Output Kits v2.5 (150 cycles) and the Illumina NextSeq500. The manufacturer’s specified parameters, combined with library loading and pooling, were used to determine the sequencing read cycles.

### scRNA-seq data processing and analysis

Raw FASTQ files of single-cell RNA sequencing data were processed using Cell Ranger 7.0.1 to generate the count matrices. Pre-built mm10, GRCh38, and barnyard reference genomes of both mouse (mm10) and human (GRCh38) downloaded from the 10X genomics website (https://support.10xgenomics.com/single-cell-geneexpression/software/downloads/latest) were used for the alignment of single-cell data from different mouse models and cell lines. scRNA-seq analysis was conducted using the R (version 4.0.0) package Seurat (version 4.0.3) (Butler et al. 2018) to process the expression matrices and perform downstream analysis. Multiple functions implemented in Seurat were used. To avoid the analysis driven by noise and low-quality cells, cells with a limited number of genes were discarded using the quantile function from R (version 4.0.0) to determine the range of genes to filter low-quality cells. The quantile range of genes was set from 2.5% to 97.5% and we discarded the cells with genes over 97.5% and less than 2.5%. Cells with more than 5% of mitochondrial counts were filtered for downstream analysis. The “Sctransform” function was used to normalize and stabilize the variance of expression matrices. The expression matrices were dimension reduced with principal component analysis (PCA). ‘FindNeighbors’ was used to define the nearest neighbors among cells in the PCA space, number of top principal components was determined by checking the plots generated by ‘JackStrawPlot’ and ‘ElbowPlot’ functions, and then ‘FindClusters’ was used to group cells with the Louvain algorithm based on the selected resolution. ‘RunUMAP’ function was used for visualizing the UMAP dimension reduction clusters. ‘DoHeatmap’ function was employed to display the top 10 genes for each meta cluster. ‘VinPlot’ function was used to show the expression probability distribution of genes across the defined cell clusters. GSEA was performed with GSEA 4.2.2 (Mootha et al., 2003; Subramanian et al., 2005) based on differentially expressed genes for each cluster and pathways with a FDR q-value < 0.1 reported.

### Flow cytometry

Tumors or pancreata were minced and digested in 1.5 mg/mL type I collagenase (Thermo Fisher Scientific) in 5 mL PBS at 37°C for 20 minutes. Subsequently, the cell pellet was resuspended in cRPMI (10% heat inactivated FBS (Atlanta Biologicals, Atlanta, Georgia, USA), 1% PS, 1 mM sodium pyruvate, and 50 μM 2-Mercaptoethanol, filtered using 70 μm strainer (Corning), and centrifuged at 600 *g* for 3 minutes at 4°C. Subsequently, cells were incubated for 5 minuutes in RBC lysis buffer (ThermoFischer). Each sample was incubated with 100 μL surface antibody cocktail diluted in FACS buffer, 20% brilliant stain buffer (BD Bioscience) and 50 μg/mL anti-mouse CD16/32 (TONBO Biosciences) for 30 minutes on ice. Fixable viability dye-efluor780 (FVD) (eBioscience) was used at 1:1000 concentration in the buffer with the antibody cocktail. Subsequently, cells were fixed with fixation buffer (BD Bioscience) and data were acquired using a BD LSR Fortessa-X20 and analysed with FlowJo v10.7.1. Antibodies panels used for flow cytometry are listed in **Table 3**.

**Table 3:** Flow cytometry antibodies.

| Antibody | Clone ID | Dilution | Vendor, catalogue number |
| --- | --- | --- | --- |
| Anti-mouse CD45-Pacific Blue | 30-F11 | 1:100 | Bio Legend, 103126 |
| Anti-mouse CD3-PE-Cy7 | 145-2C11 | 1:100 | eBioscience, 56-0032-82 |
| Anti-mouse CD4-BV605 | RM4-5 | 1:200 | BioLegend, 100548 |
| Anti-mouse CD8-BV650 | 53-6.7 | 1:200 | BioLegend, 100742 |
| Anti-mouse CD11b-BV711 | M1/70 | 1:400 | BD Bioscience, 563168 |
| Anti-mouse F4/80-PE | Cl:A3-1 | 1:50 | BioRad, MCA497PE |
| Anti-mouse CD19-PerCP-Cy5.5 | 1D3 | 1:100 | BD Bioscience, 561113 |
| Anti-mouse CD11c-PE-CF594 | HL3 | 1:100 | BD Bioscience, 562454 |
| Anti-mouse NK1.1, PE | PK136 | 1:100 | eBioscience,12-5941-83 |

### Capillary western immunoassay (WES/Simple Western)

400,000 to 500,000 cells were seeded and after 24 hours, treated with vehicle (DMSO), MRTX1133, or MRTX849 for 3 hours prior to protein isolation with RIPA buffer containing 1x cOmpleted EDTA free protease inhibitor (Roche) and 1x phosSTOP (Roche). 0.4-0.5 μg/mL of protein was prepared for Simple Western (Bio-Techne) according to manufacturer’s instructions. Samples were immunoassayed for phospho-ERK (Cell Signaling 4376S, 1:10) and vinculin (Abcam ab129002, 1:250) and anti-rabbit secondary antibody (Bio-Techne) with the 23-230 kDa separation module on a Wes instrument (30 min separation at 475 V, 5 min blocking, 30 min primary antibody, 30 min secondary antibody, 15 min chemiluminescence detection). Compass software (Bio-Techne) was used to generate gel images.

### Western blot

To analyze protein expression in MIA PaCa-2 and KPC689 cell lines and DH-50 patient derived organoids (PDOs), cell lysates were prepared in RIPA lysis buffer and protein concentrations were measured using Bicinchoninic Acid (BCA) assay (Pierce™ BCA Protein Assay Kit, Thermo Fischer). The same amount of protein was loaded for each sample (20 to 40 μg of protein lysates) and diluted with Laemmli Sample Buffer (Biorad, 1610747) and RIPA buffer in a final volume of 30 μL and incubated at 95°C for 5 min. Subsequently, the protein lysates were loaded into Bolt™ 4-12% Bis-Tris Plus gel (Thermo Fischer) for electrophoretic separation and transferred onto nitrocellulose membranes (Amersham™, Protran™). 5% non-fat milk in TBST was used as the blocking buffer. Following blocking, membranes were incubated overnight at 4°C in the following primary antibodies: Fas (EMD Millipore 05-351, 1:500), Phospho-p44/42 MAPK (Erk1/2) (Cell Signaling 4376S, 1:1000) and Vinculin (Abcam ab129002, 1:10,000). Membranes were incubated in secondary antibodies (Donkey Anti-rabbit IgG H&L: Abcam ab16284, 1:10,000 for vinculin primary and Peroxidase AffiniPure Goat Anti-Rat IgG (H+L): Jackson Immunoresearch Labs 112-035-003, 1:10,000 for Fas primary) for 1 hour at room temperature. Membranes were developed with chemiluminescence reagents (Pierce™ ECL Western Blotting Substrate) as per manufacturer’s recommendation. Uncropped, full length images from the western blots are included in the supplement.

### Chromatin immunoprecipitation

For chromatin immunoprecipitation (ChIP) experiments, 5 x 10^6^ cells per sample from KPC689 cells transfected (for 72 hours) with siScr/siKrasG12D. Two transfections, 36 hours apart were performed with Lipofectamine 2000 (Invitrogen) in OPTI-MEM (Gibco) as per the manufacturer’s instructions. Cells were trypsinized and cross-linked by gently shaking cells suspended in 1% methanol free formaldehyde (ThermoFischer) in RPMI with 10% FBS and 1% P/S at room temperature for 10-12 minutes. Subsequently, the suspension was incubated in 125 mM glycine for 5 minutes at room temperature. The cells were washed twice with ice cold PBS with 1 mM PMSF. The cell pellets were snap frozen in liquid nitrogen and ChIP assays were performed at the MD Anderson Epigenomics Profiling Core with a previously described high-throughput ChIP protocol (Blecher-Gonen et al., 2013). The resulting chromatin was incubated overnight at 4°C with histone modification antibodies (H3K27ac, Abcam ab4729; H3K27me3, Diagenode C15410195 and H3, Abcam ab1791) pre-conjugated with Dynabeads Protein G. ChIPs with non-histone antibodies (DNMT1, Novus Biologicals NB100-56519 and EZH2, Cell Signaling 5246S) were performed using the iDeal ChIP kit (Diagenode) with modifications to manufacturer instructions. The immunocomplexes were collected following day using Dynamag, washed, treated with RNase and Proteinase K, and reverse crosslinked overnight followed by DNA extraction. The DNA region of interest was detected by real-time qPCRs using following oligonucleotides at FAS transcription start site for histone enrichment; 5’-CTGCCTCTGGTAAGCTTTGG, 3’-CAGCCACATCTGGAATCTCA, and at FAS promoter for DNMT1 and EZH2 enrichment; 5’-CCCTGTATTCCCATTCATCG and 3’-ACTAGGGGAGGGGACAGAAA. The statistical analyses were performed on transcript level normalized to IgG (for DNMT1 and EZH2 ChIPs) and on transcript level normalized to histone H3 (for H3K27me3 and H3K27ac ChIPs).

### siRNA knockdown experiments and quantitative real-time PCR analyses

The siRNA sequences for siKras^G12D^ and siScr used for iExo treatment or in vitro lipofectamine transfection are described below and have been validated in previous studies (Kamerkar et al., 2017; Mendt et al., 2018). The siKras^G12D^ sequence is: sense strand 5’ -GUUGGAGCUGAUGGCGUAGTT-3’, Antisense 5’ -CUACGCCAUCAGCUCCAACTT-3’) and siScr sequence is: sense strand 5’-UUCUCCGAACGUGUCACGUTT-3’, Antisense 5’-ACGUGACACGUUCGGAGAATT-3’. RNA from transfected cells were tested for knockdown of *Fas* and *Kras^G12D^* by qPCR. Primer sequences include *ACTB:* F-5’ AGAAAATCTGGCACCACACC 3’ and R-5’ AGAGGCGTACAGGGATAGCA 3’. *FAS*: F-5’ GGACCCAGAATACCAAGTGCAG 3’ and R-5’ GTTGCTGGTGAGTGTGCATTCC 3’. qPCR was run using Fast SYBR Green Master Mix (Applied Biosystems #4385612) and the QuantStudio 7 Flex Real-Time PCR System (Applied Biosystems). The relative fold change in gene expression was determined using the 2^-ΔΔCt^ method, with the control group was arbitrarily set to 1. Statistical analyses were computed on biological replicates values of ΔCt.

### Statistical analyses

Graphical representation of the data and statistical tests were performed using GraphPad Prism 8 and reported in the respective figure legends. Shapiro-Wilk test was used to assess normality of distribution of samples. For two groups comparison, unpaired T-test (for samples with normal distribution) and Mann-Whitney test (for samples with non-normal distribution) was used for comparison of means. For nominal variables, Kruskal-Wallis test was used. For multiple groups comparison, one-way analysis of variance (ANOVA) with Tukey’s, Holm-Sidak’s or Dunnett’s multiple comparisons test (for samples with normal distribution) and Kruskal-Wallis: Dunn’s multiple comparisons test (for samples with non-normal distribution) was used. Two-way ANOVA with Tukey’s/ Sidak’s multiple comparisons test was used for comparison of histopathological analysis of tissue phenotypes. Log-rank test was used to compare Kaplan-Meier survival curves. P values are reported as *P<0.05, **P<0.01, ***P<0.001, **** P<0.0001, *ns*: not significant.

## Conflict of interest

UT MD Anderson Cancer Center and R.K. hold patents in the area of exosome biology (unrelated to the topic of this publication) and are licensed to Codiak Biosciences Inc. MD Anderson Cancer Center and R.K. are stock equity holders in Codiak Biosciences Inc. R.K. is a consultant and a scientific advisor of Codiak Biosciences Inc. A.M. receives royalties for a pancreatic cancer biomarker test from Cosmos Wisdom Biotechnology, and this financial relationship is managed and monitored by the UTMDACC Conflict of Interest Committee. A.M. is also listed as an inventor on a patent that has been licensed by Johns Hopkins University to Thrive Earlier Detection and serves as a consultant for Freenome and Tezcat Biosciences.

## Acknowledgements

This work was supported by Break Through Cancer, Cambridge, MA. KKM and KMM were supported by an Ergon Foundation Post-Doctoral Trainee Fellowship. RK is a Distinguished University Chair supported by Sid W. Richardson Foundation. The Kalluri laboratory exosomes studies are also supported in part by NCI R35 CA263815 and PDAC related studies by NCI P01 CA117969. AM is also supported by the Sheikh Khalifa bin Zayed Foundation and NCI U54CA274371. Other support includes the Small Animal Imaging Facility and veterinary pathology services core facility at MD Anderson Cancer Center supported by NCI P30CA16672. We thank the MDACC Cytogenetics and Cell Authentication Core for STR testing, Lori Wilson and Traver Hart for assistance with sequencing, Nicolas Ryujin and Shabnam Shalapour for assistance with FACS sorting, Charles Kingsley for assistance with animal imaging, Sushrut Kamerkar for assistance with iExosomes experiments, Sarah Patel for assistance with tissue processing, and Xiaoyan Ma, Andy Zuniga, Briana Epps, Diego Torres, Robert Mullinax and Angela Harris for their assistance with PDX related studies.

## Author contribution

KKM, KMM, VSL, AM, TPH and RK conceived and designed the project, and wrote or edited the manuscript. HY provided PKC-HY19636 cells. KKM, KMM, VSL, HL, SY, BL, AMS, MLK, SJM, BAMD, HS, KAA, SK, YB, NF, MPK, AML and PJK performed experiments and analyzed data. PAG generated the PDO. Specifically, HL and TPH designed, performed, and analyzed the PDX related studies. KKM, KMM, VSL, MLK designed, performed, and supervised non-PDX in vivo studies. HS, SJM, and AMS treated non-PDX mice, assisted with tissue collection, and tissue processing and section. BAM and PJK assisted with tissue processing and fibroblast staining. SY, KAA, and BL performed experiments and/or analyzed scRNA sequending studies. KMM, YB, and SY performed in vitro studies. KKM, KMM, VSL, HL, SY and BL generated figures.

## Notes

### Summary of Updates

error in citation updated

## References

Alam, A., Levanduski, E., Denz, P., Villavicencio, H.S., Bhatta, M., Alhorebi, L., Zhang, Y., Gomez, E.C., Morreale, B., Senchanthisai, S., et al. (2022). Fungal mycobiome drives IL-33 secretion and type 2 immunity in pancreatic cancer. Cancer Cell 40, 153–167 e111.

Blecher-Gonen, R., Barnett-Itzhaki, Z., Jaitin, D., Amann-Zalcenstein, D., Lara-Astiaso, D., and Amit, I. (2013). High-throughput chromatin immunoprecipitation for genome-wide mapping of in vivo protein-DNA interactions and epigenomic states. Nat Protoc 8, 539–554.

Brembeck, F.H., Schreiber, F.S., Deramaudt, T.B., Craig, L., Rhoades, B., Swain, G., Grippo, P., Stoffers, D.A., Silberg, D.G., and Rustgi, A.K. (2003). The mutant K-ras oncogene causes pancreatic periductal lymphocytic infiltration and gastric mucous neck cell hyperplasia in transgenic mice. Cancer Res 63, 2005–2009.

Canon, J., Rex, K., Saiki, A.Y., Mohr, C., Cooke, K., Bagal, D., Gaida, K., Holt, T., Knutson, C.G., Koppada, N., et al. (2019). The clinical KRAS(G12C) inhibitor AMG 510 drives anti-tumour immunity. Nature 575, 217–223.

Carugo, A., Genovese, G., Seth, S., Nezi, L., Rose, J.L., Bossi, D., Cicalese, A., Shah, P.K., Viale, A., Pettazzoni, P.F., et al. (2016). In Vivo Functional Platform Targeting Patient-Derived Xenografts Identifies WDR5-Myc Association as a Critical Determinant of Pancreatic Cancer. Cell Rep 16, 133–147.

Chang, S.H., Mirabolfathinejad, S.G., Katta, H., Cumpian, A.M., Gong, L., Caetano, M.S., Moghaddam, S.J., and Dong, C. (2014). T helper 17 cells play a critical pathogenic role in lung cancer. Proc Natl Acad Sci U S A 111, 5664–5669.

Chen, Y., McAndrews, K.M., and Kalluri, R. (2021). Clinical and therapeutic relevance of cancer-associated fibroblasts. Nat Rev Clin Oncol 18, 792–804.

Chen, Z., Trotman, L.C., Shaffer, D., Lin, H.K., Dotan, Z.A., Niki, M., Koutcher, J.A., Scher, H.I., Ludwig, T., Gerald, W., et al. (2005). Crucial role of p53-dependent cellular senescence in suppression of Pten-deficient tumorigenesis. Nature 436, 725–730.

Collins, M.A., Bednar, F., Zhang, Y., Brisset, J.C., Galban, S., Galban, C.J., Rakshit, S., Flannagan, K.S., Adsay, N.V., and Pasca di Magliano, M. (2012). Oncogenic Kras is required for both the initiation and maintenance of pancreatic cancer in mice. J Clin Invest 122, 639–653.

Commisso, C., Davidson, S.M., Soydaner-Azeloglu, R.G., Parker, S.J., Kamphorst, J.J., Hackett, S., Grabocka, E., Nofal, M., Drebin, J.A., Thompson, C.B., et al. (2013). Macropinocytosis of protein is an amino acid supply route in Ras-transformed cells. Nature 497, 633–637.

Deng, Y., Xia, X., Zhao, Y., Zhao, Z., Martinez, C., Yin, W., Yao, J., Hang, Q., Wu, W., Zhang, J., et al. (2021). Glucocorticoid receptor regulates PD-L1 and MHC-I in pancreatic cancer cells to promote immune evasion and immunotherapy resistance. Nat Commun 12, 7041.

Dey, P., Li, J., Zhang, J., Chaurasiya, S., Strom, A., Wang, H., Liao, W.T., Cavallaro, F., Denz, P., Bernard, V., et al. (2020). Oncogenic KRAS-Driven Metabolic Reprogramming in Pancreatic Cancer Cells Utilizes Cytokines from the Tumor Microenvironment. Cancer Discov 10, 608–625.

Elyada, E., Bolisetty, M., Laise, P., Flynn, W.F., Courtois, E.T., Burkhart, R.A., Teinor, J.A., Belleau, P., Biffi, G., Lucito, M.S., et al. (2019). Cross-Species Single-Cell Analysis of Pancreatic Ductal Adenocarcinoma Reveals Antigen-Presenting Cancer-Associated Fibroblasts. Cancer Discov 9, 1102–1123.

Fung-Leung, W.P., Schilham, M.W., Rahemtulla, A., Kundig, T.M., Vollenweider, M., Potter, J., van Ewijk, W., and Mak, T.W. (1991). CD8 is needed for development of cytotoxic T cells but not helper T cells. Cell 65, 443–449.

Hallin, J., Bowcut, V., Calinisan, A., Briere, D.M., Hargis, L., Engstrom, L.D., Laguer, J., Medwid, J., Vanderpool, D., Lifset, E., et al. (2022). Anti-tumor efficacy of a potent and selective non-covalent KRAS(G12D) inhibitor. Nat Med 28, 2171–2182.

Hallin, J., Engstrom, L.D., Hargis, L., Calinisan, A., Aranda, R., Briere, D.M., Sudhakar, N., Bowcut, V., Baer, B.R., Ballard, J.A., et al. (2020). The KRAS(G12C) Inhibitor MRTX849 Provides Insight toward Therapeutic Susceptibility of KRAS-Mutant Cancers in Mouse Models and Patients. Cancer Discov 10, 54–71.

Hingorani, S.R., Petricoin, E.F., Maitra, A., Rajapakse, V., King, C., Jacobetz, M.A., Ross, S., Conrads, T.P., Veenstra, T.D., Hitt, B.A., et al. (2003). Preinvasive and invasive ductal pancreatic cancer and its early detection in the mouse. Cancer Cell 4, 437–450.

Hingorani, S.R., Wang, L., Multani, A.S., Combs, C., Deramaudt, T.B., Hruban, R.H., Rustgi, A.K., Chang, S., and Tuveson, D.A. (2005). Trp53R172H and KrasG12D cooperate to promote chromosomal instability and widely metastatic pancreatic ductal adenocarcinoma in mice. Cancer Cell 7, 469–483.

Hiraoka, N., Onozato, K., Kosuge, T., and Hirohashi, S. (2006). Prevalence of FOXP3+ regulatory T cells increases during the progression of pancreatic ductal adenocarcinoma and its premalignant lesions. Clin Cancer Res 12, 5423–5434.

Huang, L., Holtzinger, A., Jagan, I., BeGora, M., Lohse, I., Ngai, N., Nostro, C., Wang, R., Muthuswamy, L.B., Crawford, H.C., et al. (2015). Ductal pancreatic cancer modeling and drug screening using human pluripotent stem cell- and patient-derived tumor organoids. Nat Med 21, 1364–1371.

Jackson, E.L., Willis, N., Mercer, K., Bronson, R.T., Crowley, D., Montoya, R., Jacks, T., and Tuveson, D.A. (2001). Analysis of lung tumor initiation and progression using conditional expression of oncogenic K-ras. Genes Dev 15, 3243–3248.

Kamerkar, S., LeBleu, V.S., Sugimoto, H., Yang, S., Ruivo, C.F., Melo, S.A., Lee, J.J., and Kalluri, R. (2017). Exosomes facilitate therapeutic targeting of oncogenic KRAS in pancreatic cancer. Nature 546, 498–503.

Kemp, S.B., Cheng, N., Markosyan, N., Sor, R., Kim, I.K., Hallin, J., Shoush, J., Quinones, L., Brown, N.V., Bassett, J.B., et al. (2023). Efficacy of a Small-Molecule Inhibitor of KrasG12D in Immunocompetent Models of Pancreatic Cancer. Cancer Discov 13, 298–311.

Lanman, B.A., Allen, J.R., Allen, J.G., Amegadzie, A.K., Ashton, K.S., Booker, S.K., Chen, J.J., Chen, N., Frohn, M.J., Goodman, G., et al. (2020). Discovery of a Covalent Inhibitor of KRAS(G12C) (AMG 510) for the Treatment of Solid Tumors. J Med Chem 63, 52–65.

McAllister, F., Bailey, J.M., Alsina, J., Nirschl, C.J., Sharma, R., Fan, H., Rattigan, Y., Roeser, J.C., Lankapalli, R.H., Zhang, H., et al. (2014). Oncogenic Kras activates a hematopoietic-to-epithelial IL-17 signaling axis in preinvasive pancreatic neoplasia. Cancer Cell 25, 621–637.

McAndrews, K.M., Chen, Y., Darpolor, J.K., Zheng, X., Yang, S., Carstens, J.L., Li, B., Wang, H., Miyake, T., Correa de Sampaio, P., et al. (2022). Identification of Functional Heterogeneity of Carcinoma-Associated Fibroblasts with Distinct IL6-Mediated Therapy Resistance in Pancreatic Cancer. Cancer Discov 12, 1580–1597.

Mendt, M., Kamerkar, S., Sugimoto, H., McAndrews, K.M., Wu, C.C., Gagea, M., Yang, S., Blanko, E.V.R., Peng, Q., Ma, X., et al. (2018). Generation and testing of clinical-grade exosomes for pancreatic cancer. JCI Insight 3.

Mootha, V.K., Lindgren, C.M., Eriksson, K.F., Subramanian, A., Sihag, S., Lehar, J., Puigserver, P., Carlsson, E., Ridderstrale, M., Laurila, E., et al. (2003). PGC-1alpha-responsive genes involved in oxidative phosphorylation are coordinately downregulated in human diabetes. Nat Genet 34, 267–273.

Olive, K.P., Tuveson, D.A., Ruhe, Z.C., Yin, B., Willis, N.A., Bronson, R.T., Crowley, D., and Jacks, T. (2004). Mutant p53 gain of function in two mouse models of Li-Fraumeni syndrome. Cell 119, 847–860.

Ostrem, J.M., Peters, U., Sos, M.L., Wells, J.A., and Shokat, K.M. (2013). K-Ras(G12C) inhibitors allosterically control GTP affinity and effector interactions. Nature 503, 548–551.

Ozdemir, B.C., Pentcheva-Hoang, T., Carstens, J.L., Zheng, X., Wu, C.C., Simpson, T.R., Laklai, H., Sugimoto, H., Kahlert, C., Novitskiy, S.V., et al. (2014). Depletion of carcinoma-associated fibroblasts and fibrosis induces immunosuppression and accelerates pancreas cancer with reduced survival. Cancer Cell 25, 719–734.

Pylayeva-Gupta, Y., Grabocka, E., and Bar-Sagi, D. (2011). RAS oncogenes: weaving a tumorigenic web. Nat Rev Cancer 11, 761–774.

Raphael, B.J., Hruban, R.H., Aguirre, A.J., Moffitt, R.A., Yeh, J.J., Stewart, C., Robertson, A.G., Cherniack, A.D., Gupta, M., Getz, G., et al. (2017). Integrated Genomic Characterization of Pancreatic Ductal Adenocarcinoma. Cancer Cell 32, 185–203.e113.

Rhim, A.D., Oberstein, P.E., Thomas, D.H., Mirek, E.T., Palermo, C.F., Sastra, S.A., Dekleva, E.N., Saunders, T., Becerra, C.P., Tattersall, I.W., et al. (2014). Stromal elements act to restrain, rather than support, pancreatic ductal adenocarcinoma. Cancer Cell 25, 735–747.

Soucek, L., Whitfield, J., Martins, C.P., Finch, A.J., Murphy, D.J., Sodir, N.M., Karnezis, A.N., Swigart, L.B., Nasi, S., and Evan, G.I. (2008). Modelling Myc inhibition as a cancer therapy. Nature 455, 679–683.

Subramanian, A., Tamayo, P., Mootha, V.K., Mukherjee, S., Ebert, B.L., Gillette, M.A., Paulovich, A., Pomeroy, S.L., Golub, T.R., Lander, E.S., et al. (2005). Gene set enrichment analysis: a knowledge-based approach for interpreting genome-wide expression profiles. Proc Natl Acad Sci U S A 102, 15545–15550.

Wang, X., Allen, S., Blake, J.F., Bowcut, V., Briere, D.M., Calinisan, A., Dahlke, J.R., Fell, J.B., Fischer, J.P., Gunn, R.J., et al. (2022). Identification of MRTX1133, a Noncovalent, Potent, and Selective KRAS(G12D) Inhibitor. J Med Chem 65, 3123–3133.

Yao, W., Rose, J.L., Wang, W., Seth, S., Jiang, H., Taguchi, A., Liu, J., Yan, L., Kapoor, A., Hou, P., et al. (2019). Syndecan 1 is a critical mediator of macropinocytosis in pancreatic cancer. Nature 568, 410–414.

Ying, H., Kimmelman, A.C., Lyssiotis, C.A., Hua, S., Chu, G.C., Fletcher-Sananikone, E., Locasale, J.W., Son, J., Zhang, H., Coloff, J.L., et al. (2012). Oncogenic Kras maintains pancreatic tumors through regulation of anabolic glucose metabolism. Cell 149, 656–670.

Zdanov, S., Mandapathil, M., Abu Eid, R., Adamson-Fadeyi, S., Wilson, W., Qian, J., Carnie, A., Tarasova, N., Mkrtichyan, M., Berzofsky, J.A., et al. (2016). Mutant KRAS Conversion of Conventional T Cells into Regulatory T Cells. Cancer Immunol Res 4, 354–365.

Zhang, Y., Velez-Delgado, A., Mathew, E., Li, D., Mendez, F.M., Flannagan, K., Rhim, A.D., Simeone, D.M., Beatty, G.L., and Pasca di Magliano, M. (2017). Myeloid cells are required for PD-1/PD-L1 checkpoint activation and the establishment of an immunosuppressive environment in pancreatic cancer. Gut 66, 124–136.

Zheng, X., Carstens, J.L., Kim, J., Scheible, M., Kaye, J., Sugimoto, H., Wu, C.C., LeBleu, V.S., and Kalluri, R. (2015). Epithelial-to-mesenchymal transition is dispensable for metastasis but induces chemoresistance in pancreatic cancer. Nature 527, 525–530.

