## Supplementary Figures for "Oncogenic Kras^G12D^ specific non-covalent inhibitor reprograms tumor microenvironment to prevent and reverse early pre-neoplastic pancreatic lesions and in combination with immunotherapy regresses advanced PDAC in a CD8^+^ T cells dependent manner"

### Supplemental Figure 1

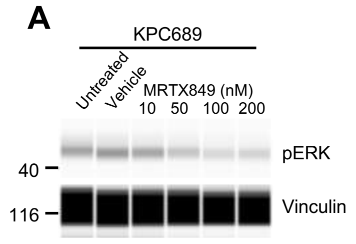


B

| Cell line | KRAS mutation status | MRTX1133 IC50 (nM) |
|---|---|---|
| HPAC | G12D | 6.076 |
| KPC689 | G12D | 47.96 |
| A549 | G12S | n.d. |
| HCT116 | G13D | 1128 |
| PSN-1 | G12R | 3641 |

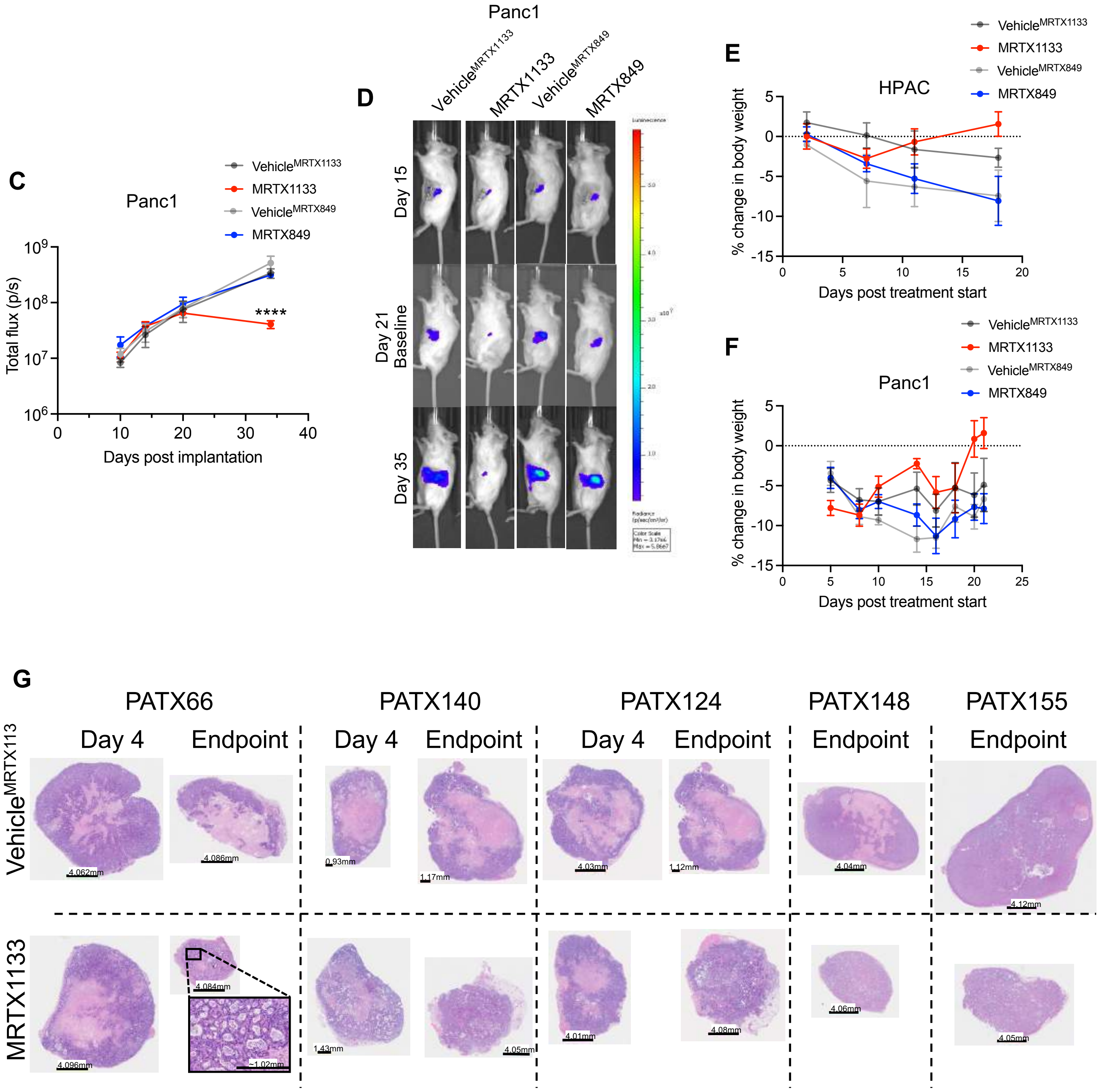

### Supplemental Figure 2

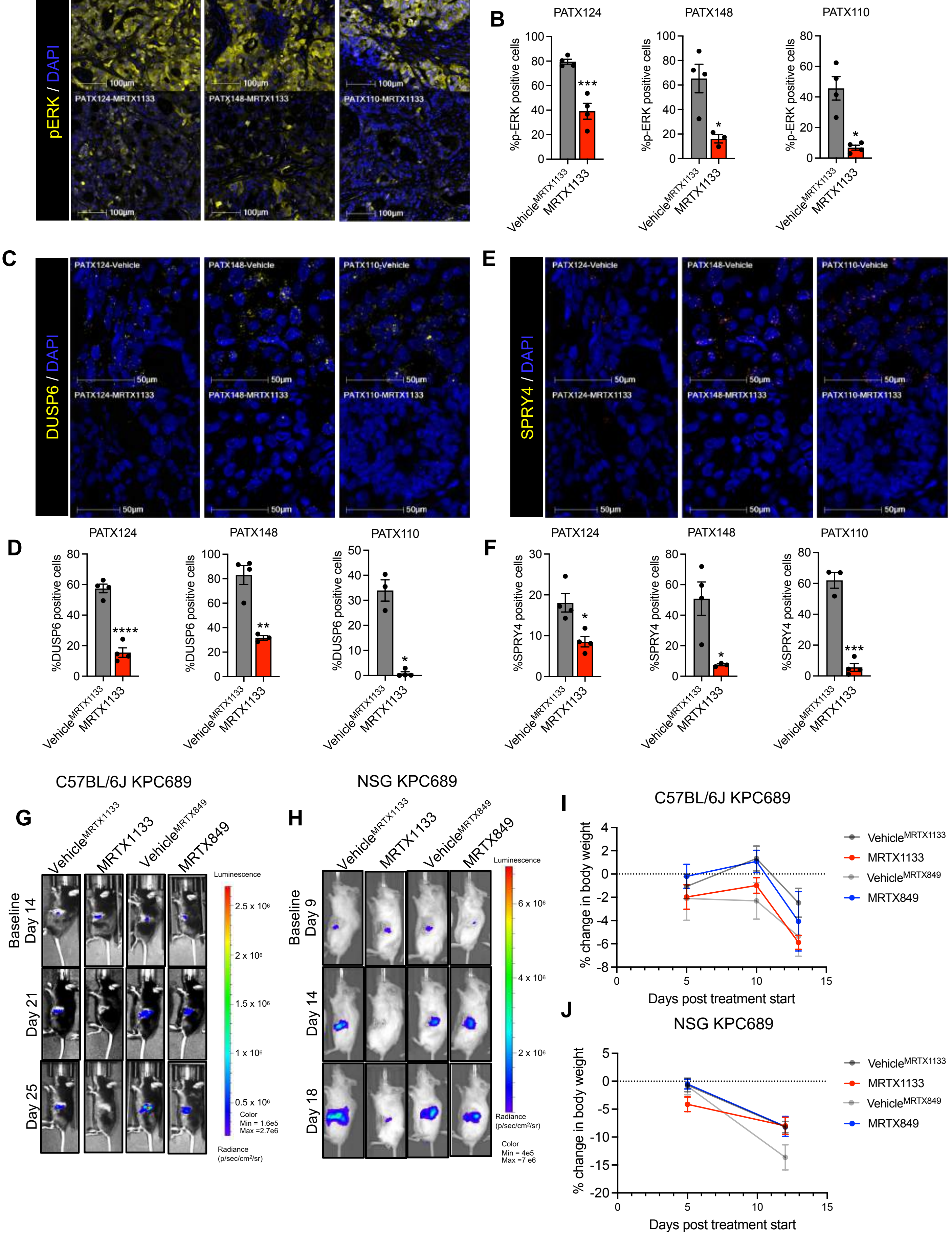

### Supplemental Figure 3

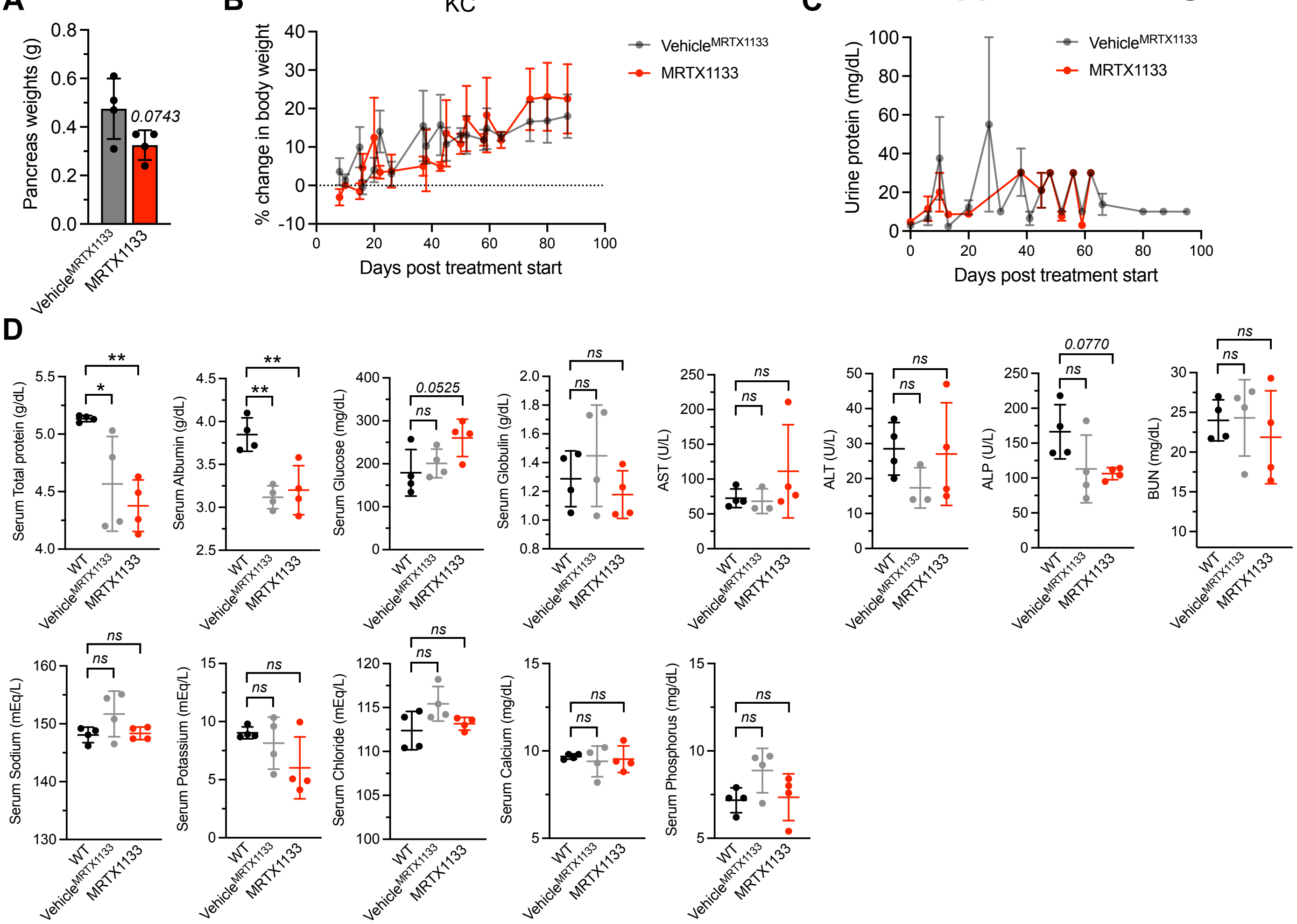


| Serum Total bilirubin (mg/dL) | | |
|---|---|---|
| C57BL6/J | KC | |
| WT | Vehicle^MRTX1133 | MRTX1133 |
| <0.15 | <0.15 | <0.15 |
| <0.15 | <0.15 | <0.15 |
| <0.15 | <0.15 | <0.15 |
| <0.15 | <0.15 | <0.15 |

| Serum Creatinine (mg/dL) | | |
|---|---|---|
| C57BL6/J | KC | |
| WT | Vehicle^MRTX1133 | MRTX1133 |
| <0.20 | <0.20 | 0.22 |
| 0.23 | <0.20 | <0.20 |
| <0.20 | 0.24 | <0.20 |
| 0.24 | <0.20 | <0.20 |

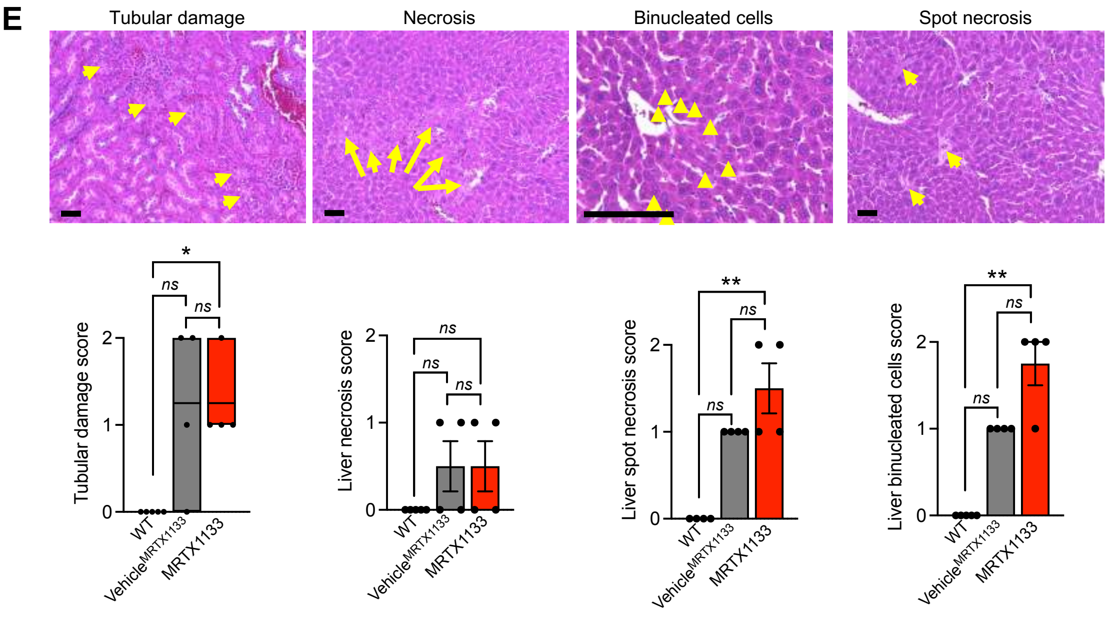

### Supplemental Figure 4

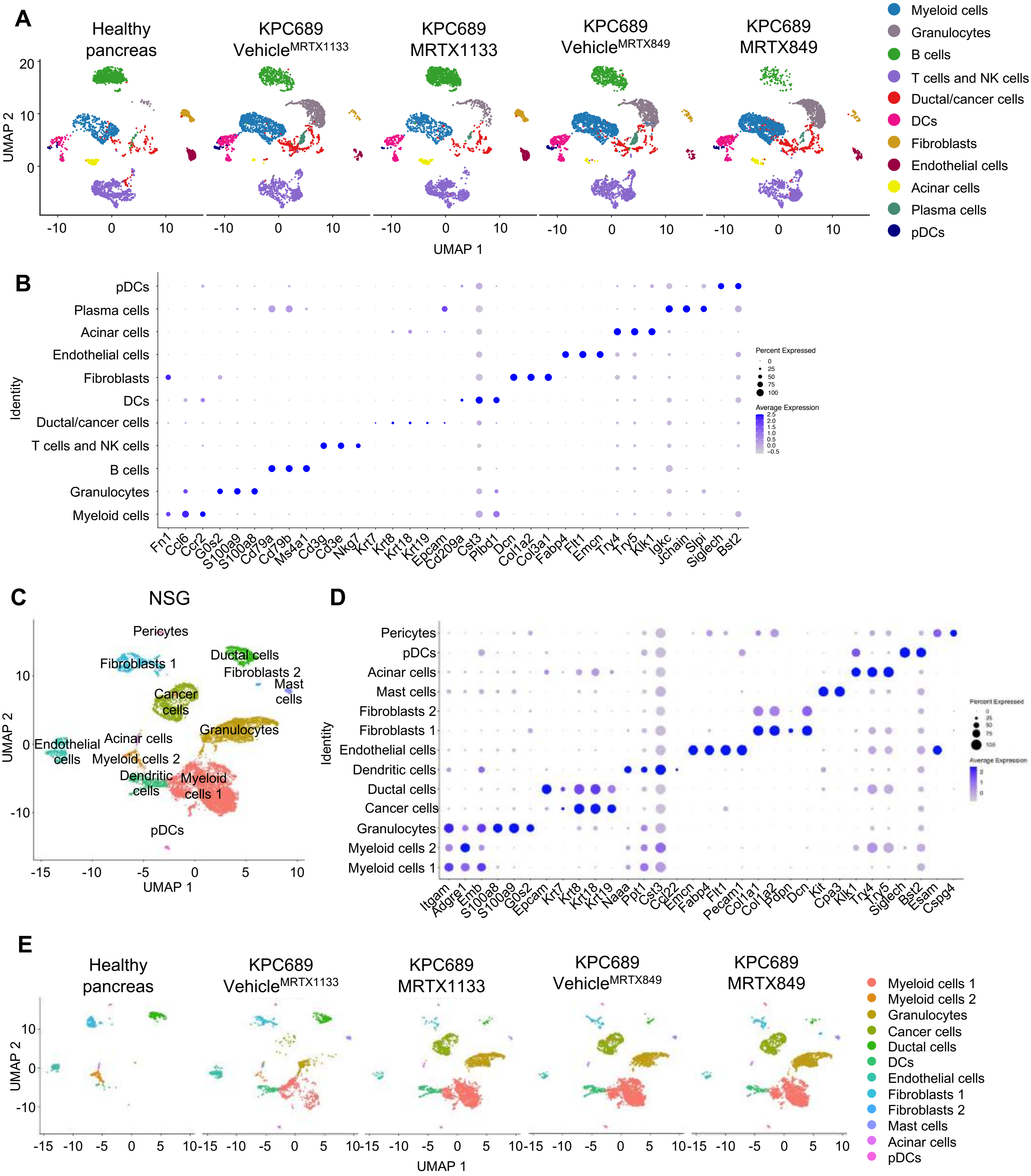

### Supplemental Figure 5

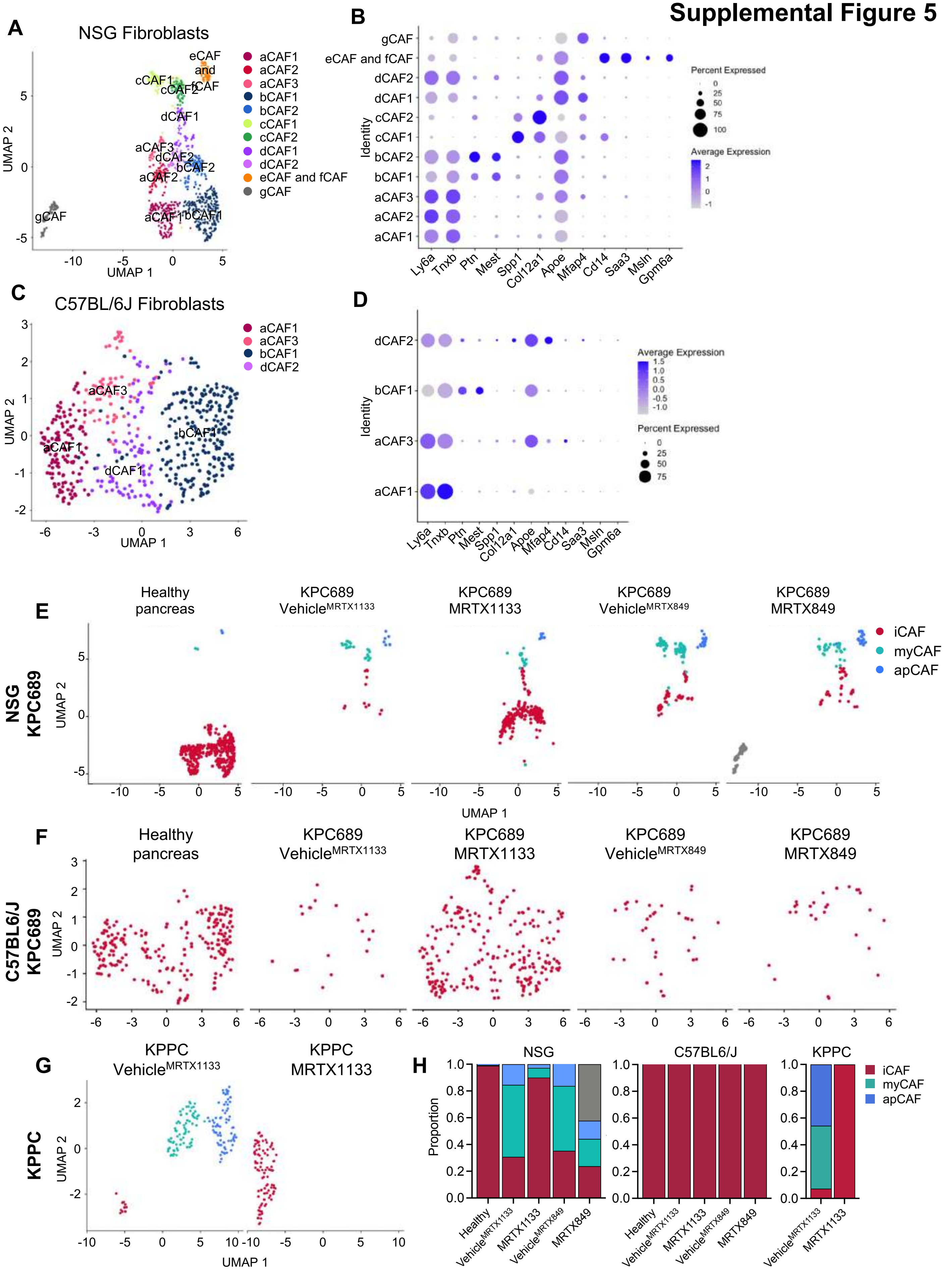

### Supplemental Figure 6

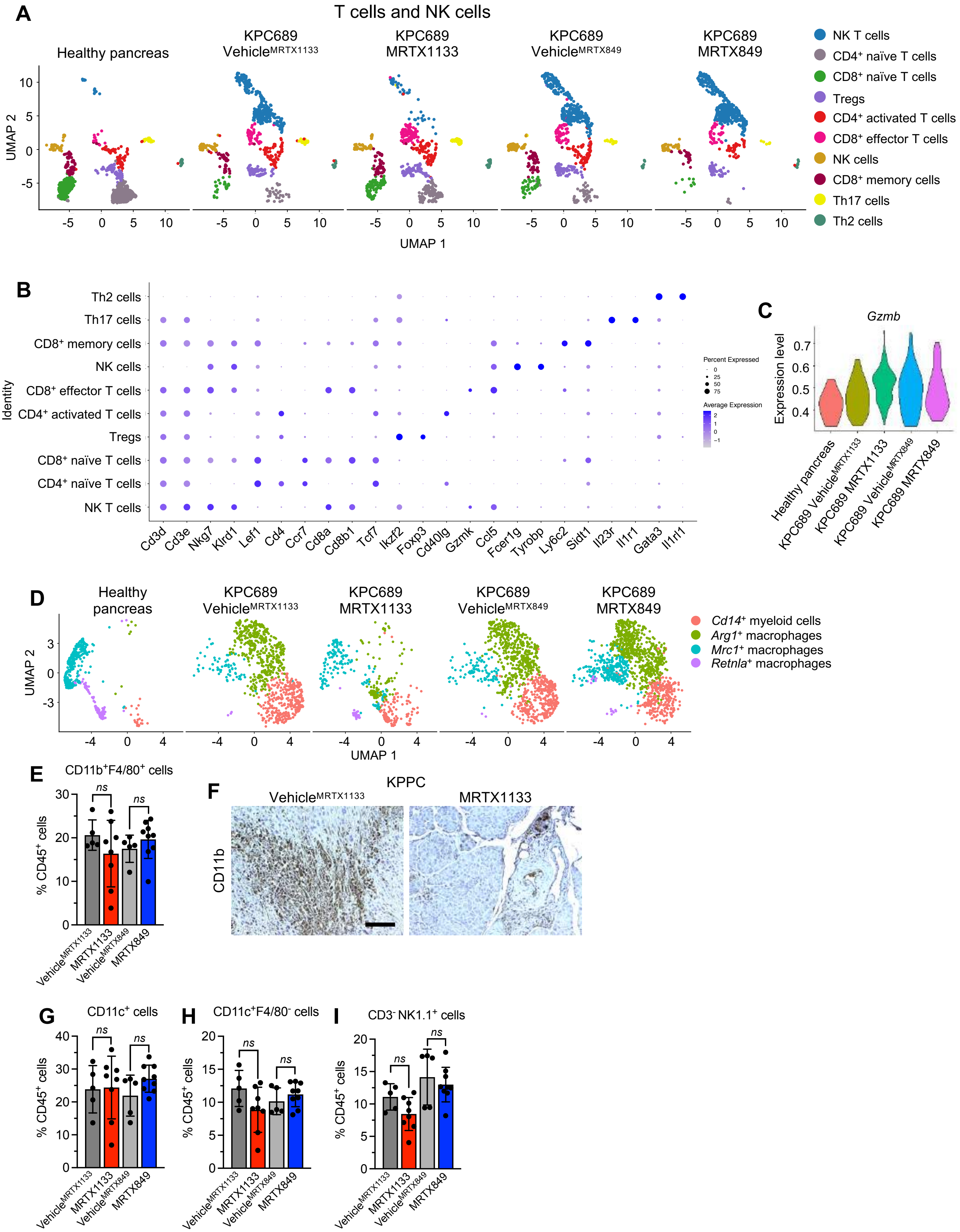

### Supplemental Figure 7

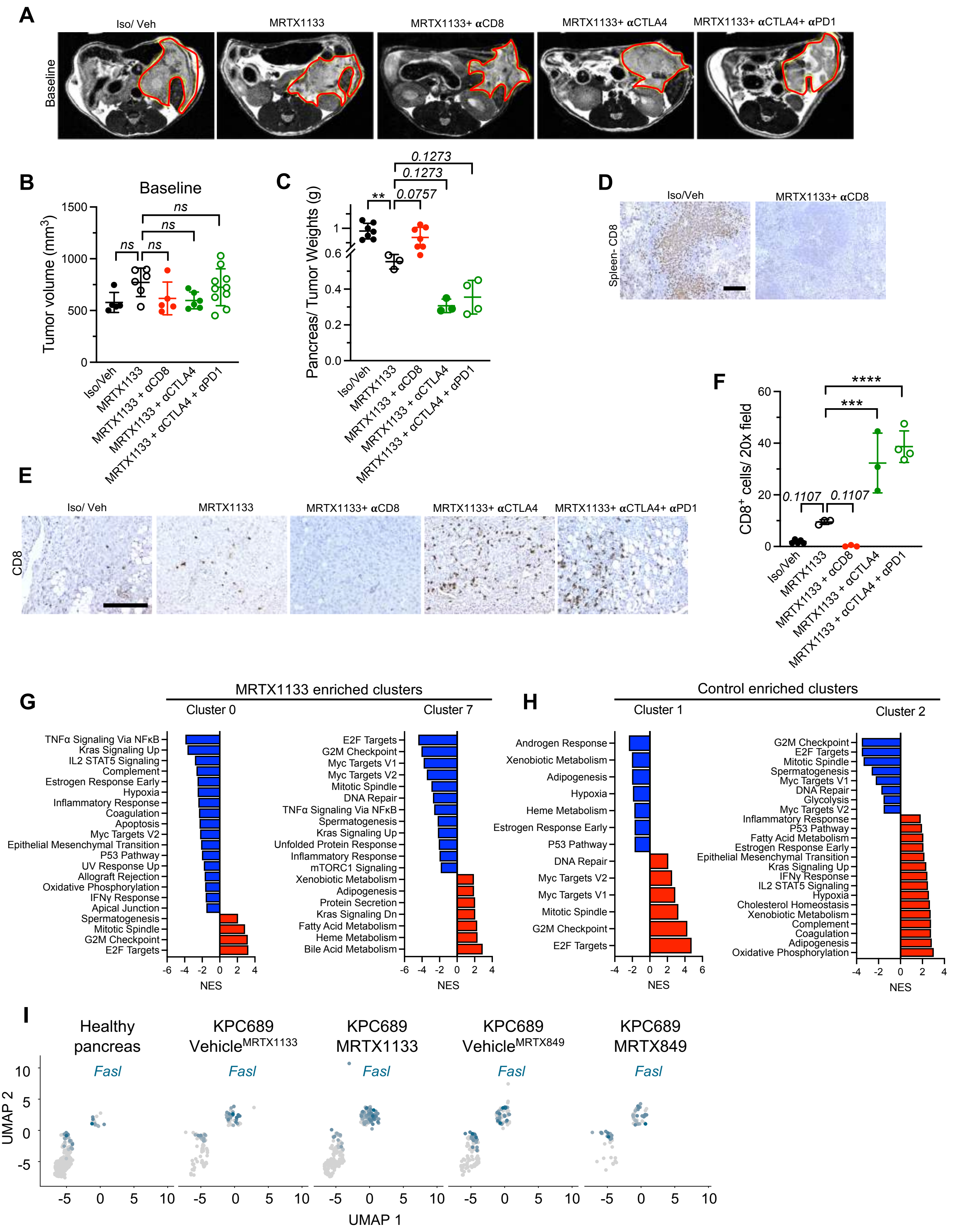

### Supplemental Figure 8

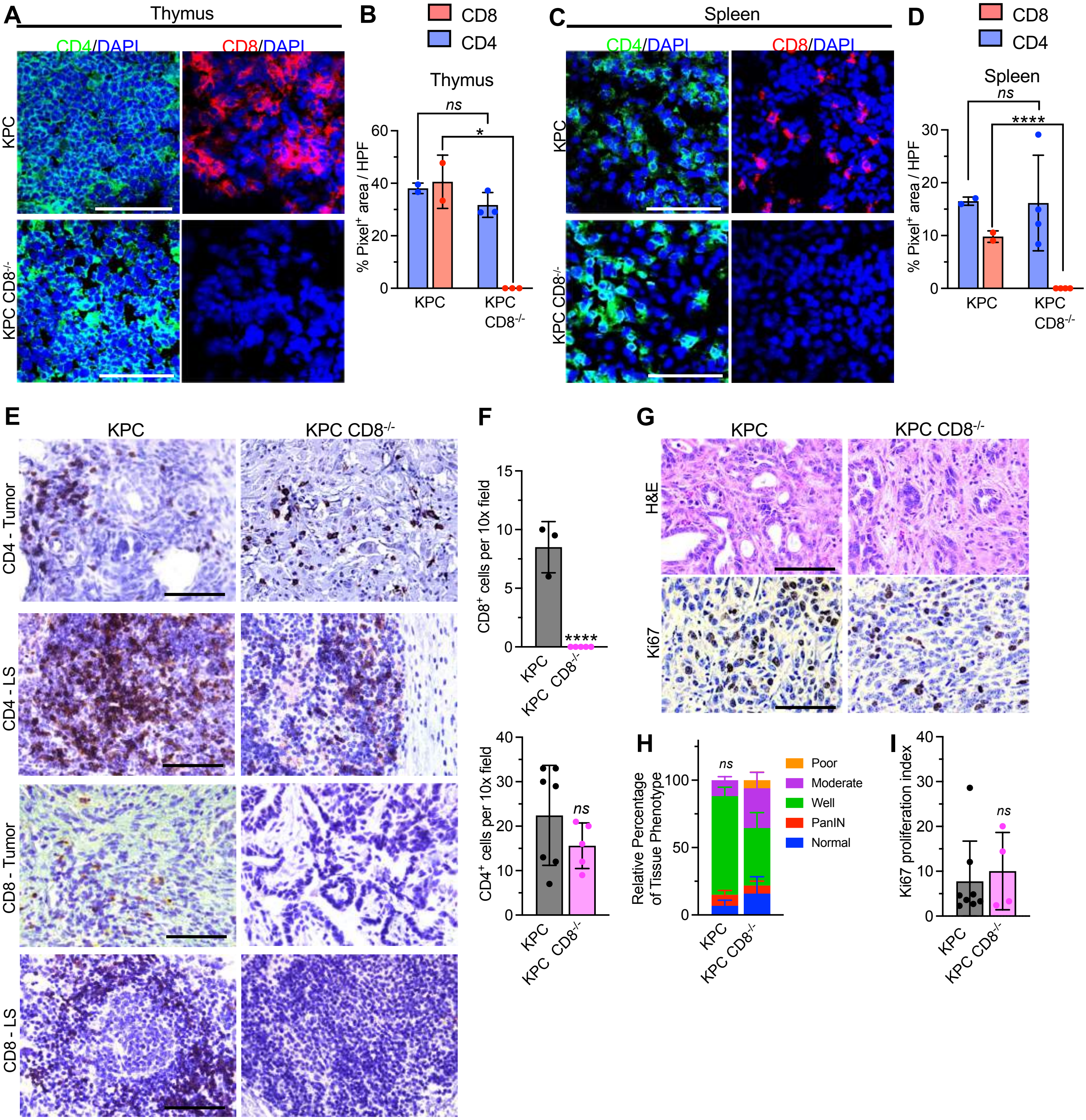

**Uncropped blots**

Figure 1A

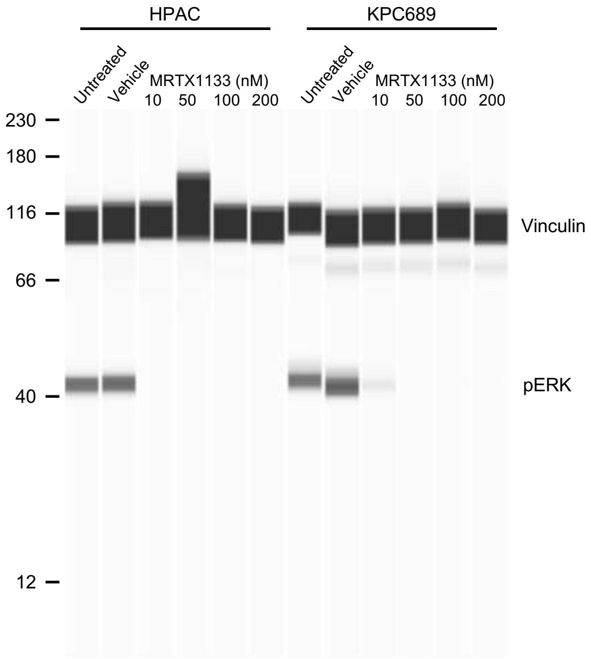


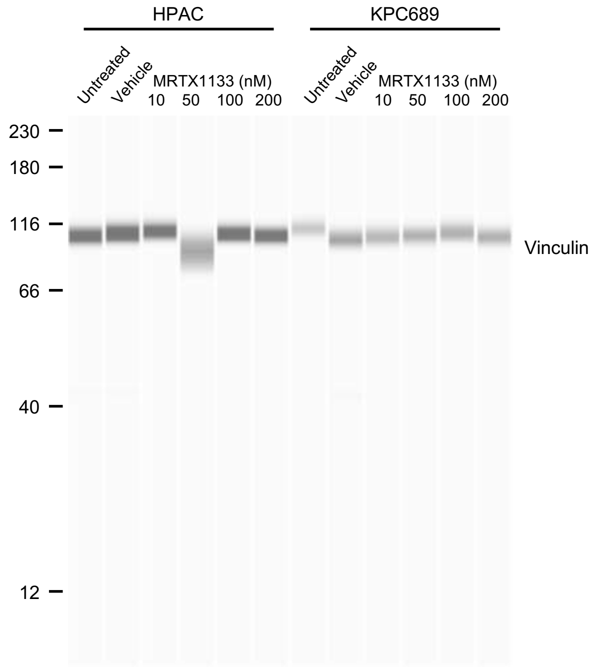


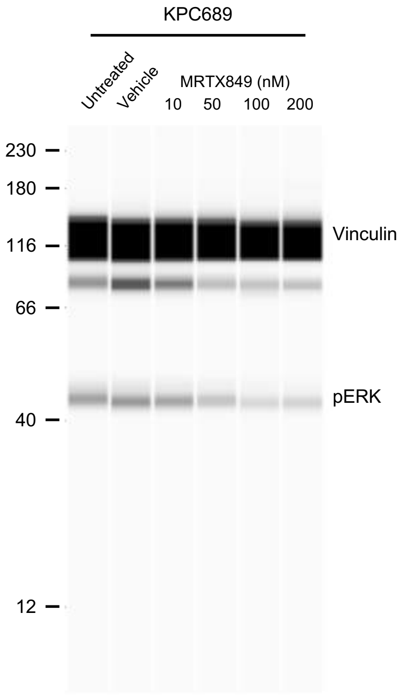

**Uncropped blots**

#### Figure 6E

p-ERK
AZD6244 (MEKi)
Miapaca-2

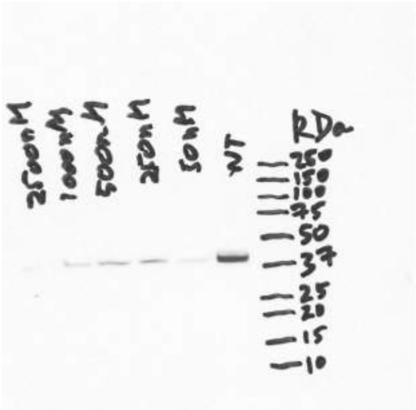


Vinculin
AZD6244 (MEKi)
Miapaca-2

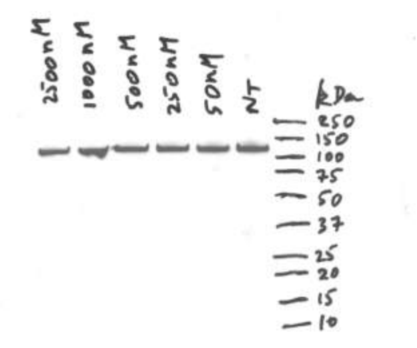


#### Figure 6G

p-ERK
MRTX1133
KPC-689

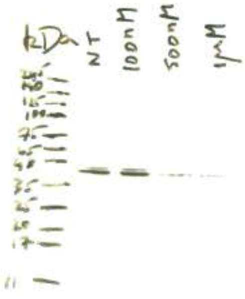


Fas
MRTX1133
DH-50

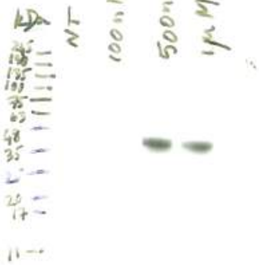


Vinculin
MRTX1133
DH-50

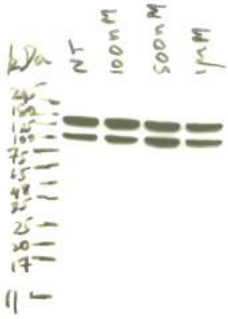

**Uncropped blots**

Figure 6H

p-ERK
MRTX849
MIA Paca-2

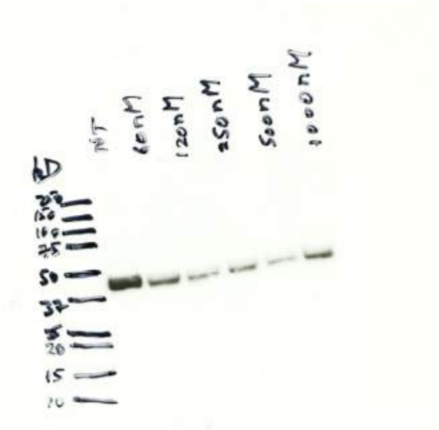

Vinculin
MRTX849
MIA Paca-2

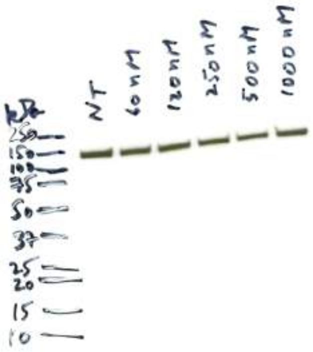

Figure 6K

p-ERK
MRTX849
DH-50

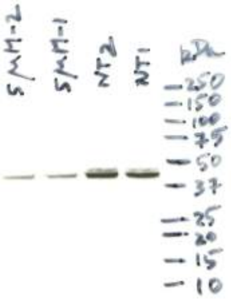

Vinculin
MRTX849
DH-50

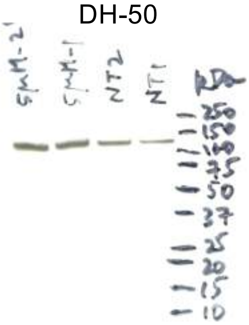

Figure 6N

p-ERK
MRTX1133
DH-50

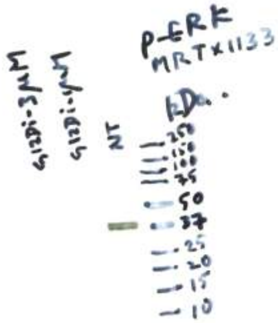

Vinculin
MRTX1133
DH-50

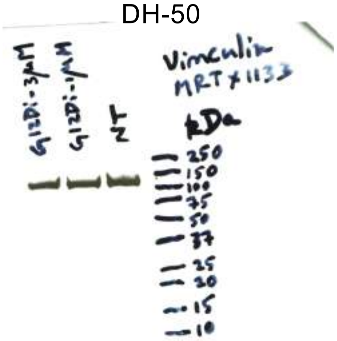
